# Temporal regulation of epithelium formation

**DOI:** 10.1101/076570

**Authors:** Stephen E. Von Stetina, Jennifer Liang, Georgios Marnellos, Susan E. Mango

**Affiliations:** Department of Molecular and Cellular Biology, BL3015, 16 Divinity Avenue, Cambridge, MA 02138; Informatics and Scientific Applications, Science Division, Faculty of Arts and Sciences, Harvard University,MA 02138, Cambridge USA

**Keywords:** *par-3*/ASIP, *dlg-1*/Discs Large and *erm-1*/Ezrin Radixin Moesin, breast cancer, lung cancer, centralspindlin, PHA-4/FOXA

## Abstract

To establish the animal body plan, embryos link the external epidermis to the internal digestive tract. In *Caenorhabditis elegans,* this linkage is achieved by the Arcade Cells, which form an epithelial bridge between the foregut and epidermis, but little is known about how development of these three epithelia is coordinated temporally. The Arcade Cell epithelium is generated after the epidermis and digestive tract epithelia have matured, ensuring that both organs can withstand the mechanical stress of embryo elongation; mis-timing of epithelium formation leads to defects in morphogenesis. Here, we report that temporal regulation of the Arcade Cell epithelium is mediated by the pioneer transcription factor PHA-4/FoxA, the cytoskeletal regulator ZEN-4/MKLP and the polarity protein PAR-6. We find that PHA-4 activates expression of a broad cohort of epithelial genes. However, accumulation of protein is delayed by ZEN-4, acting in concert with its partner CYK-4/MgcRacGAP. Finally, PAR-6 localizes factors within adherens junctions and at the apical surface, leading to Arcade Cell polarity. The results reveal that the timing of a landmark event during embryonic morphogenesis is mediated by the concerted action of four proteins that delay the formation of an epithelial bridge. In addition, we find that FoxA associates with many epithelial genes in mammals, suggesting that regulation of epithelial identity may be a conserved feature of FoxA factors and a contributor to FoxA function in development and cancer.

## INTRODUCTION

Animal embryos coordinate the morphogenesis of their body and internal organs in a process that requires epithelia. Communication between mesenchyme and epithelia is necessary for morphogenesis of the mammalian kidney and lung, for example, as well as the *Drosophila* trachea (reviewed in (Hogan and Kolodziej, 2002; Caviglia and Luschnig, 2014; Combes *et al.*, 2015; McCulley *et al.*, 2015)). There is also synergy between different organs that helps shape the embryonic body plan. In *C. elegans*, the embryo elongates four-fold to generate a long, thin worm from an oblong ball of cells (reviewed in (Chisholm and Hardin, 2005; Vuong-Brender *et al.*, 2016)). Elongation of the body is coordinated with that of the gut by physically linking the outer epidermis to the inner digestive tract. At the anterior, the link is constructed by the Arcade Cells, which generate an epithelial bridge between the epidermis and foregut just prior to body elongation. Proper morphogenesis depends critically on this linkage. Disruption of Arcade Cell polarization by mutation or laser ablation blocks attachment of the gut tube to the epidermis (Heid *et al.*, 2001; Portereiko and Mango, 2001; Portereiko *et al.*, 2004; Mango, 2009; Kuzmanov *et al.*, 2014). As a consequence, the foregut fails to elongate, and affected animals cannot feed (Portereiko *et al.*, 2004). Similarly, in *sma-1* mutants, the foregut attaches to the epidermis but the embryo body fails to extend fully, and this leads to defects in foregut morphogenesis (McKeown *et al.*, 1998). These phenotypes indicate that proper gut morphogenesis requires attachment to the elongating epidermis. Conversely, if the Arcade Cells generate an epithelial bridge prior to epidermal maturation, the digestive tract distorts the morphology of the nose and produces feeding defects (Kelley *et al.*, 2015). These observations illustrate that embryos rely on epithelia to coordinate body morphogenesis, and raise the question of how cells dictate the timing of epithelium formation and attachment.

Epithelia become polarized with distinct apical and basolateral domains, separated by a junctional domain (Nelson *et al.*, 2013; Roignot *et al.*, 2013). Following cell-contact mediated cues, three major polarity complexes are recruited to the membrane of fly and vertebrate epithelia, and partition the cell into apical and basolateral domains (reviewed in (St Johnston and Ahringer, 2010; Roignot *et al.*, 2013; Rodriguez-Boulan and Macara, 2014)). The Crumbs complex along with the PAR complex are important for apical identity and/or junction formation (Wodarz *et al.*, 1995, 1993; Gao *et al.*, 2002; Hurd *et al.*, 2003; Roh *et al.*, 2003; Harris and Peifer, 2004; Lemmers *et al.*, 2004; Chen and Macara, 2005; St Johnston and Ahringer, 2010; Roignot *et al.*, 2013; Rodriguez-Boulan and Macara, 2014). The basolateral surface is marked by the Scribble complex, which contains Scribble (Scrib), Disks large (DLG), and Lethal Giant Larvae (LGL) (St Johnston and Ahringer, 2010; Roignot *et al.*, 2013). Diffusion between the apical and basolateral domains is blocked by the junctional domain, which also provides tissue integrity via cell-cell adhesion (Hartsock and Nelson, 2008). The combined actions of these polarity modules establish and maintain polarity in a range of tissues.

In the nematode, all epithelial cells form via MET from unpolarized precursors (Portereiko *et al.*, 2004; Achilleos *et al.*, 2010). *C. elegans* epithelia have a single electron-dense junction, termed the *C. elegans* apical junction (CeAJ) (McMahon *et al.*, 2001), which has properties of both tight and adherens junctions (Pásti and Labouesse, 2014). The apicobasal distribution of factors, including those in the Crumbs, Par, and Scribble groups, is similar between worm and vertebrate epithelia (Pásti and Labouesse, 2014). The three Crumbs homologs localize to the apical surface (Bossinger *et al.*, 2001; Segbert *et al.*, 2004; Waaijers *et al.*, 2015), PAR-6 localizes to the apical domain (Leung *et al.*, 1999; McMahon *et al.*, 2001; Nance *et al.*, 2003), and DLG-1 is found in the basal half of the CeAJ (Bossinger *et al.*, 2001; Firestein and Rongo, 2001; Koppen *et al.*, 2001; McMahon *et al.*, 2001; Segbert *et al.*, 2004). Distinct epithelia rely on the same cohort of epithelially expressed genes, indicating that temporal control cannot be explained by distinct, paralogous factors acting in different organs. For example, *par-6* and *dlg-1* are critical for the integrity of adherens junctions in all tested organs, and mutants for either gene fail to elongate their body or digestive tract (Bossinger *et al.*, 2001; Firestein and Rongo, 2001; Koppen *et al.*, 2001; McMahon *et al.*, 2001; Totong *et al.*, 2007).

Transcription factors can drive both mesenchymal-to-epithelial transition (MET) and the reverse process, epithelial-to-mesenchymal transition (EMT) (Chaffer *et al.*, 2007; Lamouille *et al.*, 2014). Targets of EMT include E-cadherin and integrin(Batlle *et al.*, 2000; Cano *et al.*, 2000; Comijn *et al.*, 2001; Perez-Moreno *et al.*, 2001; Hajra *et al.*, 2002; Bolós *et al.*, 2003; Yang *et al.*, 2004, 2009; Eger *et al.*, 2005), but the direct transcription factor targets that mediate MET are largely unknown (Chaffer *et al.*, 2007). In this study, we focus on FoxA factors, which promote the development of several epithelial organs and which are often found in tumors derived from epithelia (Friedman and Kaestner, 2006). For example, mammalian FoxA1 and FoxA2 are expressed in the lung, thyroid, kidney, and pancreas, and FoxA1 is also found in the mammary gland (Besnard *et al.*, 2004; Friedman and Kaestner, 2006). *C. elegans* has one FoxA factor, *pha-4*, that functions in two epithelial organs, the gonad and foregut (Mango *et al.*, 1994; Updike and Mango, 2007). In mammals, reduction of FoxA can promote EMT in lung, liver, prostate and breast by reducing expression of Slug, E-cadherin or MMP-9 (Tang *et al.*, 2011; Wang *et al.*, 2014; Zhang *et al.*, 2015). However, homologues of Slug, E-cadherin and metalloproteases play no known role in polarizing epithelia in worms (Costa *et al.*, 1998; Raich *et al.*, 1999; Fraser *et al.*, 2000; Kamath *et al.*, 2003; Simmer *et al.*, 2003; Rual *et al.*, 2004; Sönnichsen *et al.*, 2005; Von Stetina and Mango, 2015), indicating that either FoxA factors have additional, conserved target genes, or that FoxA factors regulate distinct epithelial genes in different organs and/or species. Here we define direct targets of *C. elegans* FoxA and explore their orthologues in vertebrates.

In this work, we examine *C. elegans* epithelium formation during morphogenesis, when a ball of cells is transformed into a long, thin worm. We find that epithelia are generated just prior to the onset of their associated morphogenetic event. We focus on the Arcade Cells, which form an epithelium that bridges the epidermis and foregut during late embryogenesis. A core set of epithelial factors is activated by the pioneer factor PHA-4/FoxA, but protein accumulation and localization are delayed by ZEN- 4/MKLP, CYK-4/MgcRacGAP and PAR-6. We extend these results to FoxA factors in mammalian tissues and determine that vertebrate FoxA factors bind many orthologous target genes. The results reveal how the exquisite timing of embryonic morphogenesis depends on temporally coordinated regulation of a common core of epithelial factors at the RNA and protein level.

## RESULTS

### Overview of *C. elegans* epithelium formation

Timing of embryo development can be tracked by the number of E (endodermal) cells and by embryo shape (Figure 1; (Sulston *et al.*, 1983)). The epidermis is the first epithelium to polarize, during the late 8E stage (~200 cells (Sulston *et al.*, 1983; Podbilewicz and White, 1994; Leung *et al.*, 1999; McMahon *et al.*, 2001; Chisholm and Hardin, 2005)). This period occurs soon after the epidermal precursors are born between the 2E and mid-8E stage, and prior to epiboly at the bean stage (Sulston *et al.*, 1983). Once the epithelium has formed, it migrates over the lateral and ventral surfaces of the embryo, thereby enclosing the embryo in preparation for body elongation (Priess and Hirsh, 1986; Williams-Masson *et al.*, 1997) (Figure 1).

**Figure 1.**
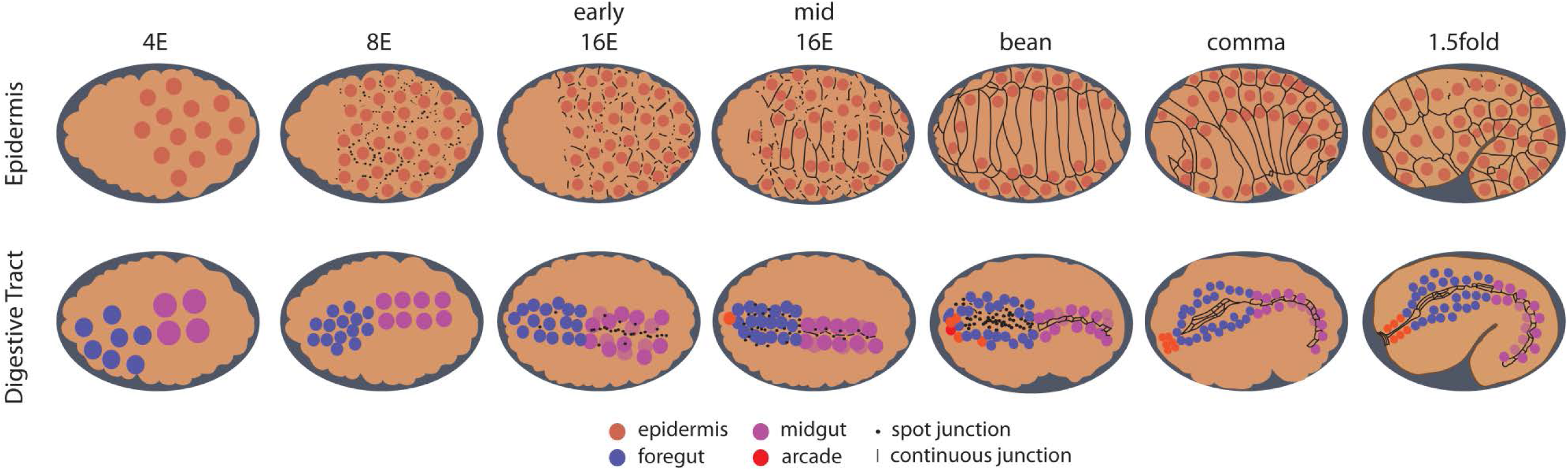
*C. elegans* embryonic stages and epithelial cell morphology. Anterior is to the left. The top row of embryos focuses on the epidermis while the bottom row depicts the digestive tract. The nuclei of the epidermis (orange), foregut (blue), midgut (magenta) and Arcade Cells (red) are shown. The staging is initially determined by the number of midgut (endoderm or E) cells that are present, but once morphogenesis begins at the bean stage the stage name refers the shape of the embryo. Junctional proteins (black) first become apparent in the epidermis in the late 8E stage, where they clump together to form spot junctions (black dots) in the 16E that resolve into continuous junctions (black lines) during the 16E stage. By the 1.5-fold stage some epidermal cells fuse, creating large multi-nucleate cells. The digestive track polarizes in a posterior to anterior direction, with the midgut expressing junctional proteins at the early 16E stage followed shortly thereafter by the foregut at the mid 16E stage. The midgut makes the transition from spot junctions to continuous junctions by the bean stage, and the foregut by the comma stage. The nine arcade cells are born around the mid-16E stage in the anterior of the developing foregut (only a subset are shown for clarity). These cells cluster together anterior to the foregut by the comma stage but do not express junctional protein until they polarize between the comma and 1.5-fold stages.

The digestive tract polarizes progressively, with midgut epithelialization commencing at the 8E-16E, and the foregut following during the mid-16E (Figure 1) (Totong *et al.*, 2007; Achilleos *et al.*, 2010). Between the comma and 1.75-fold stages, the anteriorly located Arcade Cells and posterior hindgut undergo a mesenchymal-to-epithelial transition to generate two small epithelia that link the epidermis to the foregut and hindgut respectively. The result is a torus with the epidermis on the outside and the digestive tract on the inside, held under tension by its distal attachments. The torus elongates four-fold to generate a worm with an extended, linear digestive tract (Chisholm and Hardin, 2005; Nance *et al.*, 2005). Without mechanical coupling of the gut to the epidermis, the digestive tract cannot produce its proper elongated form (Portereiko and Mango, 2001; Mango, 2009; Kelley *et al.*, 2015).

### Asynchronous accumulation of *dlg-1* RNA and protein in different organs

To understand the temporal regulation of epithelium formation, we determined the onset of expression for polarity factors by surveying members of the Par (*par-6*) and Scribble (*dlg-1*) modules with single-molecule fluorescent *in situ* hybridization (smFISH) for RNA (Ji and van Oudenaarden, 2012). PAR-6 plays an early and key role in maturing apical junctions within the epidermis and digestive tract (Totong *et al.*, 2007), whereas DLG-1 is one of the last components added to epithelial junctions and essential for their maturation (Totong *et al.*, 2007). We found that *par-6* RNA was contributed maternally, as predicted from prior studies (Watts *et al.*, 1996; Nance *et al.*, 2003), and no increase due to zygotic *par-6* RNA was detected (Figure S1; (Totong *et al.*, 2007). *dlg-1* was induced zygotically, with RNA accumulating in different organs at different times, prior to the generation of each epithelium. Thus, *dlg-1* provided a useful readout for the timing of epithelium formation.

The epidermis was the first epithelium to express *dlg-1* mRNA. It was initially detected at the late 4E stage, but with no detectable DLG-1 protein (Figure 1, Figure 2A). The level of *dlg-1* mRNA increased during the 8E stage (Figure 2B) and was maintained throughout the 16E and elongation stages (comma, 1.5fold, Figure 2C-F). DLG-1 protein was first observed during the late 8E stage, with puncta of protein visible on the membrane of nascent epidermal cells (Figure 2B). These puncta began to coalesce at the early 16E stage (Figure 2C) and formed a continuous, circumferential junction by the mid-16E stage (Figure 2D). The level of DLG-1 increased during the elongation stages (comma, 1.5-fold, Figure 2E,F), as the cells changed shape to convert the embryo from a ball into a vermiform.

**Figure 2.**
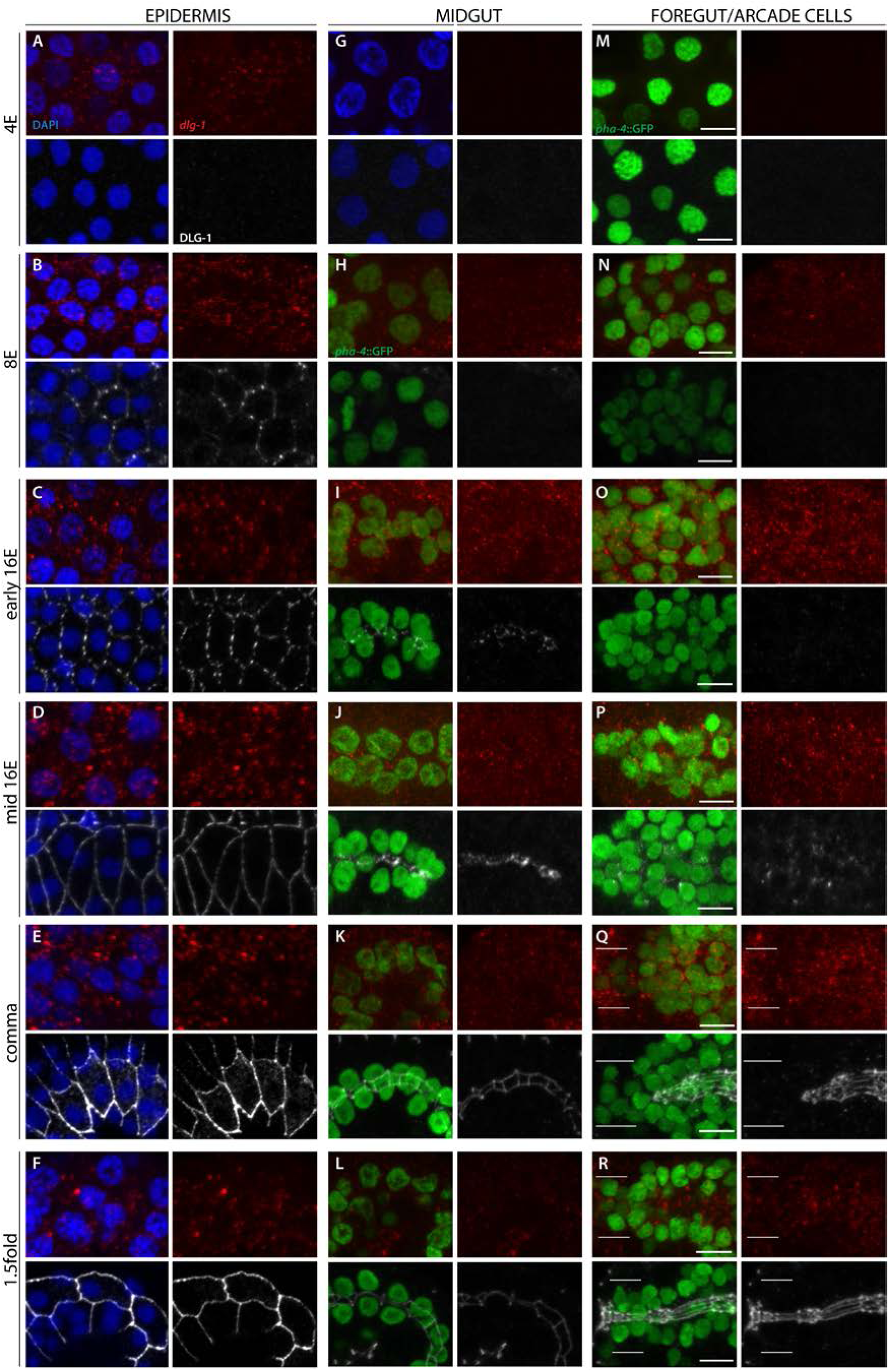
The onset of *dlg-1* RNA and protein expression in epithelia. *dlg-1* RNA is labeled in red, DLG-1 protein in white. The nuclei of the epidermis (A-F) and midgut (G) are visualized by DAPI (blue). *pha-4*::GFP (green) demarks the midgut (H-L), foregut (M-R) and Arcade Cells (white bars in Q, R). A-F. *dlg-1* RNA is first detected in the 4E epidermis (A), while the onset of protein is at the 8E stage (B). RNA and protein are continuously expressed throughout the 16E and morphogenetic stages (C-F). Note that the protein first forms spot junctions (B, C) that resolve into continuous bands (D-F). G-L. The midgut first expresses RNA at the 8E stage and protein at the early 16E. The protein first forms spot junctions (I) that quickly resolve into continuous bands (J). RNA and protein are also detected during morphogenesis (K, L). M-R. RNA expression in the foregut begins at the 8E stage (N). Unlike the midgut, protein expression is first visible as spots by the mid, rather than the early 16E, stage (compare I vs. O and P). Foregut cells form continuous junctions by the comma stage (Q). The arcade cells (white bars) have RNA but no protein at the comma stage (Q), and between the comma and 1.5fold the Arcade Cells express protein and form continuous junctions (R). All images are maximum intensity projections. Anterior is to the left. Scale bars 5 µm.

The digestive tract began to express *dlg-1* mRNA at the 8E stage (Figure 2H, N), and similar to the epidermis, the levels of RNA increased throughout the 8E and 16E stages (Figure 2I-J, O-P). DLG-1 protein was first observed in midgut precursors at the early 16E stage (Figure 2I), where puncta of protein appeared at the lateral surface and rapidly coalesced at the apical surface, in agreement with previous studies (Figure S1; (Leung *et al.*, 1999; Totong *et al.*, 2007; Achilleos *et al.*, 2010)). By the mid-16E stage (~20-40m later), the puncta of DLG-1 had banded together to form cell junctions (Figure 2J), which continued to expand and mature as the embryo elongated (comma, 1.5-fold stages, Figure 1, Figure 2K,L). The RNA remained expressed in the intestine throughout all of these stages (Figure 2I-L).

In the foregut, DLG-1 protein was first detectable by the mid-16E stage (Figure 2P), suggesting that translation of *dlg-1* mRNA was delayed in this tissue by ~20-40 minutes. We observed membrane-associated DLG-1 puncta on cell surfaces throughout the foregut at the 16E stage (Figure S1). These spots accumulated at the nascent apical surface by the bean stage (Figure S1), where they joined together to form connected junctions by the comma stage (Figure 1, Figure 2Q). The RNA remained expressed throughout these stages (Figure 2O-R). Together, these data show that the digestive tract forms in a piecemeal manner, with the midgut polarizing and maturing prior to the foregut, and both after the epidermis.

By the end of the comma stage, the epithelial cells of the foregut and midgut were linked together via adherens junctions, yet the digestive tract remained isolated from the epidermis. The digestive tract became attached to the epidermis at the anterior via epithelialization of the Arcade Cells (Figure 1) (Portereiko and Mango, 2001). The Arcade Cells are born during the mid-16E stage, starting ~290 minutes after the first division (Sulston *et al.*, 1983). The majority of these cells were found anterior to the foregut primordium and expressed *dlg-1* mRNA from birth (Figure 2O). DLG-1 protein accumulated approximately 100 minutes later in the Arcades (Figure 2R), after the epidermis and foregut had both formed epithelia and shortly before the Arcade Cells become an epithelium (i.e. between the comma and 1.25-fold stage, ~390-400 minutes after the first division)(Portereiko and Mango, 2001; Portereiko *et al.*, 2004). The presence of RNA but lack of protein was detectable by the 16E stage (Figure S2C) but clearest at the comma stage, when the Arcade Cells clustered together as a group anterior to the foregut epithelium (Figure 2Q). Thus, there was a delay in protein accumulation, suggesting that the RNA was either translationally-repressed, or that protein was made but degraded immediately. These results reveal that *dlg-1* RNA and protein are uncoupled in different organs, with the protein appearing just prior to polarity establishment.

### PHA-4 activates *dlg-1* in the foregut and Arcade Cells

The onset of *dlg-1* RNA expression suggested a transcriptional component to temporal regulation of epithelium formation. One central regulator of the digestive tract is PHA-4, a pioneer transcription factor that is also the selector gene for the *C. elegans* foregut (Horner *et al.*, 1998; Gaudet and Mango, 2002; Mango, 2009; Hsu *et al.*, 2015). To determine if PHA-4 was important for *dlg-1* expression, we performed RNAi against *pha-4* and observed reduced DLG-1 in the foregut and Arcade cells, but normal expression in the midgut and epidermis (Figure 3A). Antibody staining against endogenous PHA-4 confirmed that RNAi was effective (Figure. 3B). Apical factors PAR-3 and ERM-1 were similarly disrupted by loss of *pha-4* (Figure. 3C-F), suggesting a widespread disruption of epithelial factors.

**Figure 3.**
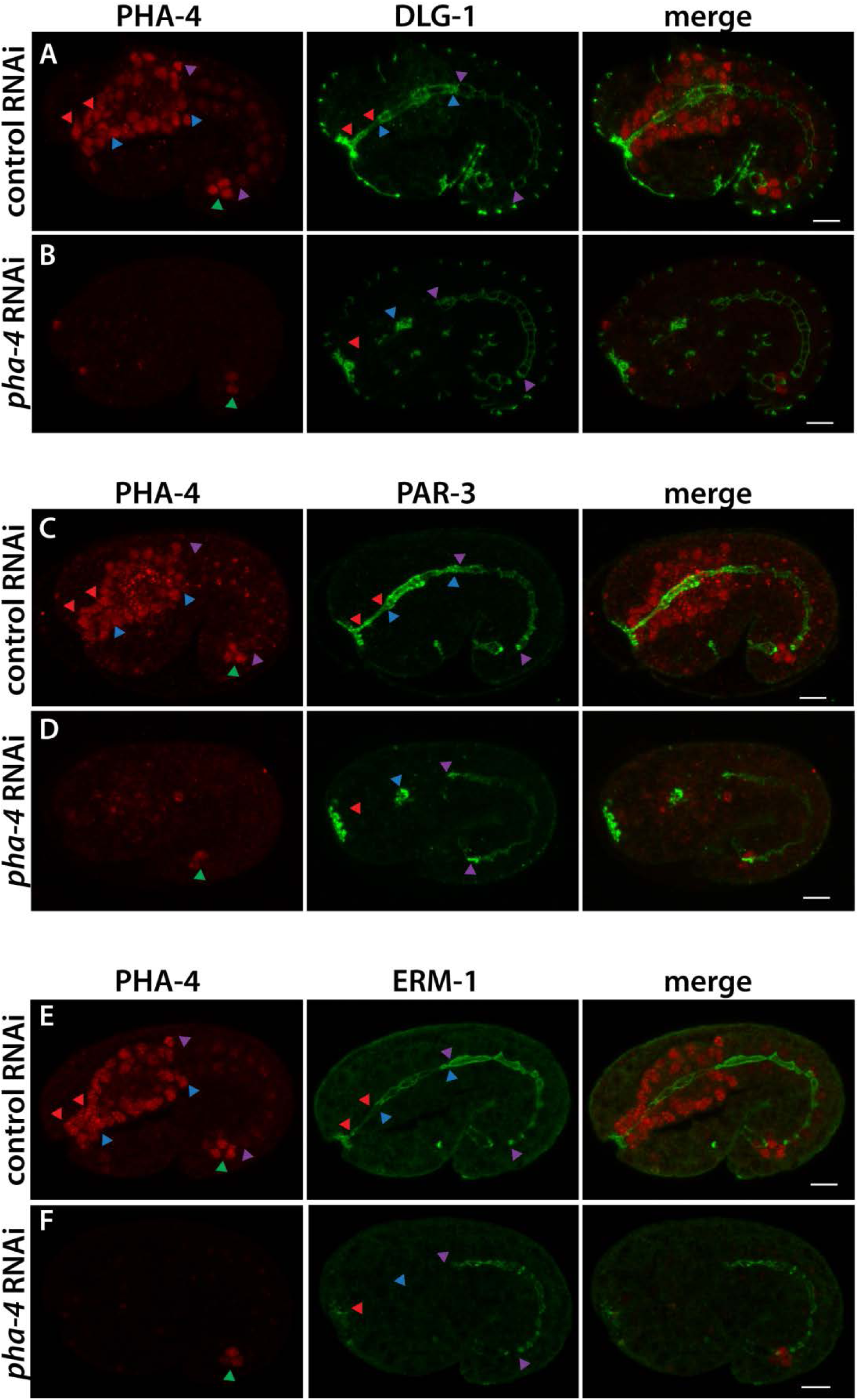
PHA-4 is required for polarity protein expression in the foregut. The Arcade Cell epithelium lies between red eads, the foregut epithelium by blue arrowheads, the midgut epithelium by purple arrowheads, and hindgut cells are d by the green arrowhead. A, C, E. In control RNAi embryos PHA-4 (red) is strongly expressed in Arcade, foregut, and hindgut nd dimly expressed in the midgut. Markers of polarity (green), such as DLG-1 (A), PAR-3 (C) and ERM-1 (E) are detected at ical or junctional domains of the digestive tract tissues. B, D, F. After *pha-4* RNAi treatment PHA-4 is undetectable in the, foregut or midgut cells. The hindgut is resistant to *pha-4(RNAi*) and serves as an internal control for antibody staining. In *pha-4(RNAi)* embryos, polarity protein expression is ablated in arcade cells (red arrowhead) and diminished in the foregut. If protein remains, it is detected in the foregut remnant (blue arrowhead). Polarity factors are unaffected in the midgut (purple arrowheads). All images are maximum intensity projections through the digestive tract. Anterior is to the left. Scale bars 5 µm.

Next we determined whether regulation of *dlg-1* transcription by PHA-4 was direct. PHA-4 binding has been mapped by ChIP-Seq (Chromatin ImmunoPrecipitation-Sequencing) (Zhong *et al.*, 2010). Examination of ChIP-seq data demonstrated two binding sites for PHA-4, one near the upstream *dlg-1* Transcription Start Site (TSS; Figure 4A; (Saito *et al.*, 2013)) and a second site just prior to the ATG start and within the first intron (Figure 4A). To test whether these sites contributed to *dlg-1* expression, we used a DLG-1::GFP reporter *P_7_dlg-1,* with 7kb of upstream sequences, which was able to rescue *dlg-1* mutants (McMahon *et al.*, 2001). This reporter expressed DLG-1 appropriately: endogenous *dlg-1* and transgenic *gfp* RNA were both detectable in the foregut and midgut at the 8E stage (i.e. at the onset of endogenous gene expression, Figure 2H and Figure S3) and abundant in these tissues by the 16E stage (Figure 4B). Live imaging demonstrated that DLG-1::GFP protein was enriched at the midline of the midgut and clustered into junctions at the 16E stage (Figure 4C). In the mid-16E, DLG-1::GFP was detectable as spots spread throughout the foregut, similar to the endogenous protein (compare Figure 4C to Figure 2P). At the comma stage, both endogenous and transgenic RNAs were detectable throughout the digestive tract (including the Arcade Cells, Figure 4K), whereas DLG-1::GFP protein, like endogenous DLG-1, was only visible in the foregut and midgut (Figure 4L). By the 1.5fold stage, DLG-1::GFP was also visible in the Arcade Cells, as expected (Figure 4U). Thus, the *P*_*7*_*dlg-1* transgene contained all the information necessary for appropriate temporal expression of RNA and protein in the digestive tract and epidermis (Figure 4D, M, V).

**Figure 4.**
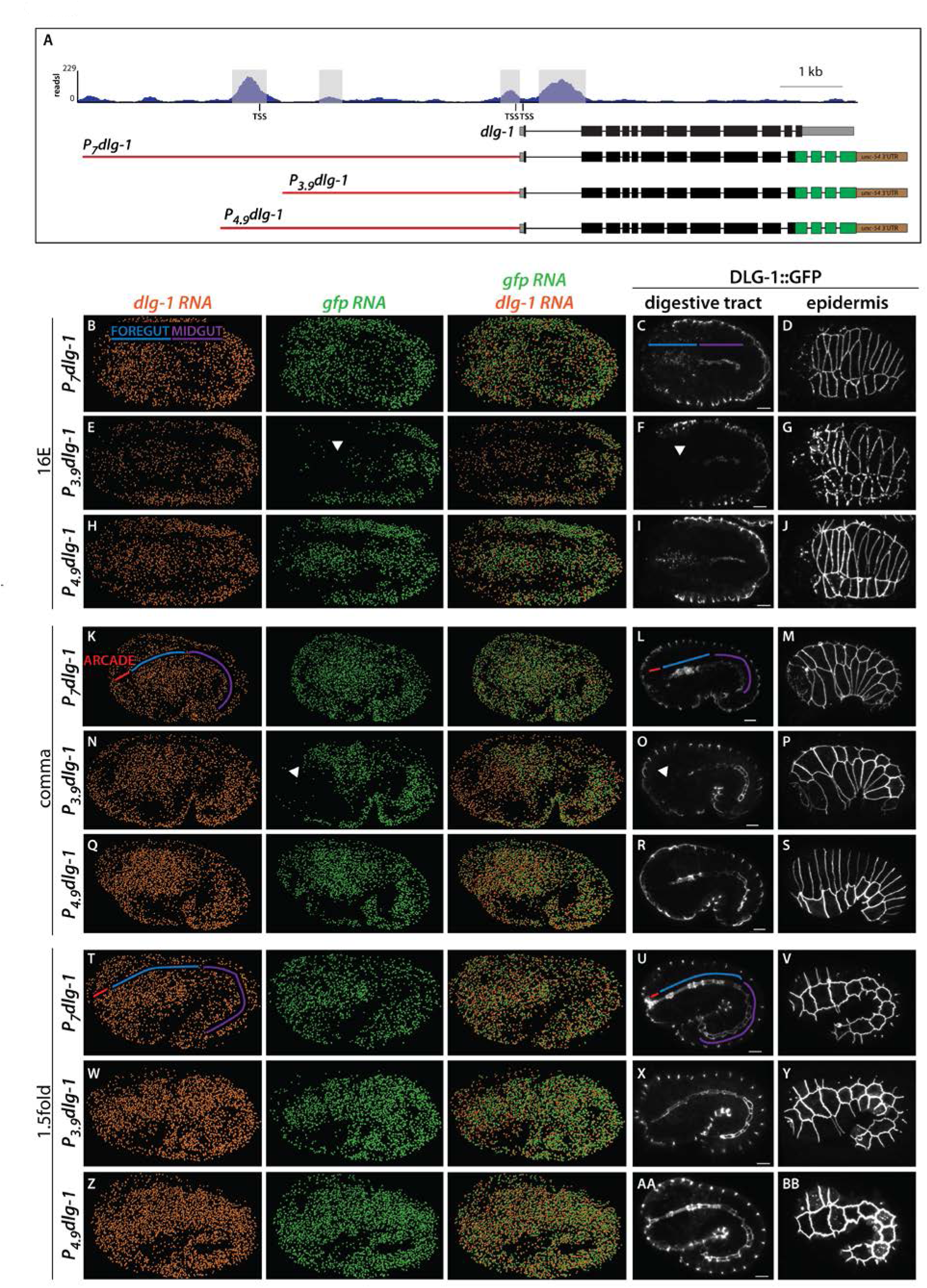
PHA-4 binding site is necessary for normal onset of *dlg-1* expression in the foregut and Arcade Cells. A. PHA-4::GFP ChIP-Seq profile at the endogenous *dlg-1* locus (top; (Zhong *et al.*, 2010)). Exons are black boxes, 5′ and 3′ UTR sequence are gray boxes, and introns are black lines. There are four called PHA-4 ChIP peaks (highlighted in gray), two surrounding the first exon, one ~3kb upstream and one ~4kb upstream, near the upstream transcription start site (TSS). Three GFP-tagged constructs are also shown, with GFP sequences in green and the heterologous *unc-54* 3’UTR in brown. The various promoter elements are shown in red. The top construct (*P*_*7*_*dlg-1;* (McMahon *et al.*, 2001)) contains 7kb of sequence, the middle (*P*_*3.9*_*dlg-1*) contains 3.9kb and excludes the most upstream PHA-4 peak, and the bottom (*P_4.9_dlg-1*) contains 4.9kb and includes the most upstream peak. B-Z. Maximum intensity projections of worm embryos expressing the various *dlg-1* promoter constructs at either the 16E, comma or 1.5fold stage. *dlg-1* mRNA detected by smFISH is pseudo-colored orange (Methods), *gfp* mRNA is pseudo-colored green, and DLG-1::GFP protein is white. The areas corresponding to the Arcade Cells are labeled with a red line, the foregut with a blue line, and the midgut with a purple line. B, K, T. Embryos expressing DLG-1::GFP under the *P*_*7*_*dlg-1* promoter have transgenic *gfp* mRNA expression that matches endogenous *dlg-1* mRNA (compare orange and green mRNA) in the Arcades, foregut, midgut and epidermis. DLG-1::GFP expression (C, D, L, M, U, V) also is similar to endogenous (compare to Figure 2). E, N, W. Embryos expressing DLG-1::GFP under the *P*_*3.9*_*dlg-1* promoter, which lacks the upstream PHA-4 peak, have transgenic *gfp* mRNA expression that matches endogenous *dlg-1* mRNA in the epidermis and midgut. Expression in the foregut and Arcade cells is delayed. In the 16E stage (E) *gfp* mRNA is nearly absent in the foregut and Arcade cells (white arrowhead). By the comma stage (N) the posterior foregut has *gfp* mRNA but the anterior foregut and Arcade Cells lack expression (white arrowhead). Transgenic mRNA expression in the Arcades and anterior foregut is seen by the 1.5fold stage (W). DLG-1::GFP protein (F, O, X) also follows this pattern (white arrowheads point to absent expression in foregut and Arcades). H, Q, Z. Embryos expressing DLG-1::GFP under the *P*_*4.9*_*dlg-1* promoter, which contains the upstream PHA-4 peak, have transgenic mRNA and protein expression profiles that match their endogenous counterparts in the Arcade Cells, foregut and epidermis. Some midgut cells show precocious gfp RNA at the 8E stage (data not shown). All images are maximum intensity projections. Anterior is to the left and scale bars are 5 µm.

We performed promoter deletion analysis to determine whether PHA-4-bound sites contributed to *dlg-1* activation. We generated worms expressing two different DLG-1::GFP constructs, one that contained all PHA-4 binding sites (*P*_*4.9*_*dlg-1 for 4.9kb*) and one that carried the site near the ATG start but lacked the major binding site at the upstream TSS (*P*_*3.9*_*dlg-1 for 3.9kb*)(Figure 3A). *P_3.9_dlg-1* faithfully recapitulated the onset of expression in the epidermis (Figure 4G, P, Y; data not shown) and midgut (Figure 4 E, F, N, O, W, X), similar to endogenous *dlg-1*. However, *P*_*3.9*_*dlg-1* was silent at the 16E stage in the foregut and Arcades, when endogenous *dlg-1* was abundant (Figure 4E, compare *gfp* RNA (green) to *dlg-1* RNA (orange)); DLG-1::GFP protein was undetectable (Figure 4F). When occasional transgenic mRNA or protein was observed, it was limited to the posterior end of the foregut (Figure S4, data not shown). Expression in the foregut of *P*_*3.9*_*dlg-1* transgenic embryos commenced slowly, progressing from posterior to anterior. Specifically, by the comma stage most of the foregut (but not the Arcade Cells) had abundant *gfp* RNA (Figure 4N), but detectable DLG-1::GFP expression was restricted to the posterior foregut (Figure 4O). At the 1.5fold stage, transgenic mRNA and protein were detected throughout the digestive tract, and in most cases included the Arcade Cells (Figure 4W, X). Adding back 1kb of sequence (*P_4.9_dlg-1*), including the upstream PHA-4 binding site, restored wild-type expression in the foregut (Figure 4H, I, Q, R, Z, AA). These data suggest that the PHA-4 binding site near the upstream TSS is critical for onset of expression in the foregut, whereas other PHA-4 binding sites and sites for other factors mediate late foregut expression (or maintenance of expression), and expression in the midgut and epidermis.

In summary, three lines of evidence suggest PHA-4 contributes to the proper onset of *dlg-1* expression: First, DLG-1 expression was reduced or absent from *pha-4* mutant foreguts and Arcade cells. Second, PHA-4 ChIP peaks were detected within *dlg-1* regulatory elements (Zhong *et al.*, 2010), and third, deletion analysis implicated the upstream PHA-4 binding site (~4kb upstream of the ATG start) for normal onset of expression in the foregut and Arcade Cells (but not the midgut or epidermis). Thus, distinct regulatory elements control the onset of *dlg-1* expression in foregut and Arcade Cells vs. midgut and epidermis.

### PHA-4 activates an epithelial program during foregut polarization

To extend our observations beyond *dlg-1*, we examined PHA-4 association with genes encoding other polarity proteins, relying again on the modENCODE PHA-4 ChIP-seq dataset (Zhong *et al.*, 2010). Strikingly, genes for every epithelial polarity protein annotated as expressed in the embryonic foregut (27/27) had an associated PHA-4 peak, many close to the transcription start site (TSS) or within the first large intron, which are common sites for *bona fide* PHA-4 targets (Gaudet and Mango, 2002) (Figure 5, Table 1). For example, *par-3* had two PHA-4 ChIP peaks within the large, first intron (which is a promoter for the epithelial-specific isoform (Achilleos *et al.*, 2010)), *erm-1* had an upstream peak and one in a large intron, and the α-catenin *hmp-1* contained one peak at the TSS. In support of PHA-4 regulating these epithelial-specific genes, we found reduced levels of ERM-1 and PAR-3 in *pha-4(RNAi)* mutant foreguts (Figure 3C-F).

**Figure 5.**
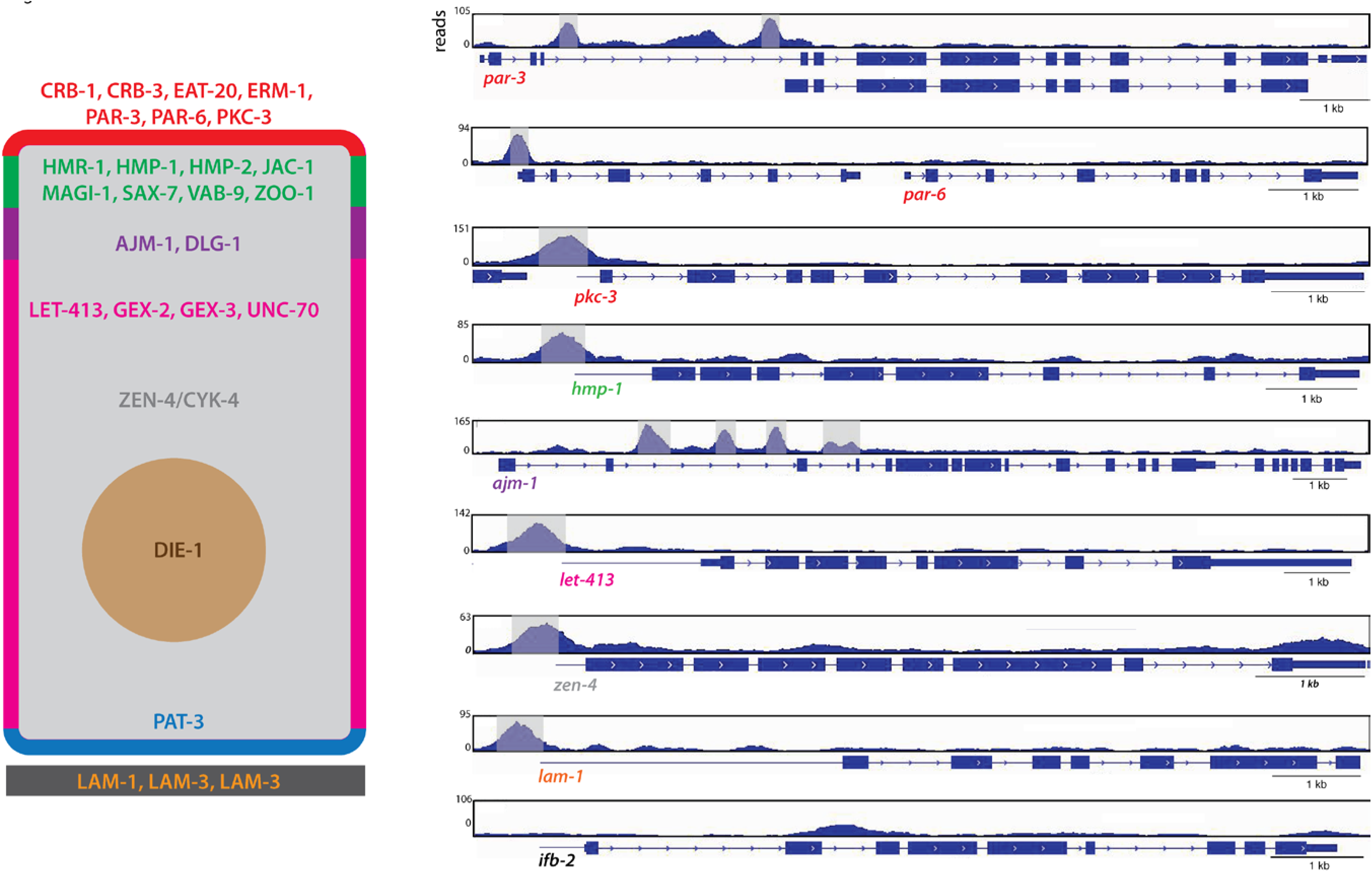
PHA-4 binds to the regulatory elements of epithelial factors. The left cartoon represents a generic *C. elegans* epithelial cell, with the apical domain led red, apical junction green, basal junction purple, lateral domain fuschia, basal domain blue, extracellular matrix black, cytoplasm gray and nucleus n. A subset of epithelial factors whose regulatory regions are bound by PHA-4 (Table 1) are listed in their respective domains. The genome browser ls on the right display the PHA-4 binding peaks for selected genes, adapted from Zhong et al, 2010. Exons are denoted by blue boxes and introns by blue with arrowheads. A blue line extends from the annotated transcription start site (Kruesi *et al.*, 2013) to the first exon for genes (such as *hmp-1*) where TSS) is >100bp upstream of the first exon. Significant PHA-4 peaks are highlighted in gray. Note that the smaller isoform of *par-3* is epithelial specific, esting that the large upstream intron in which PHA-4 binds could regulate the expression of this isoform specifically. *par-6* is the second gene in an

**Table 1.**
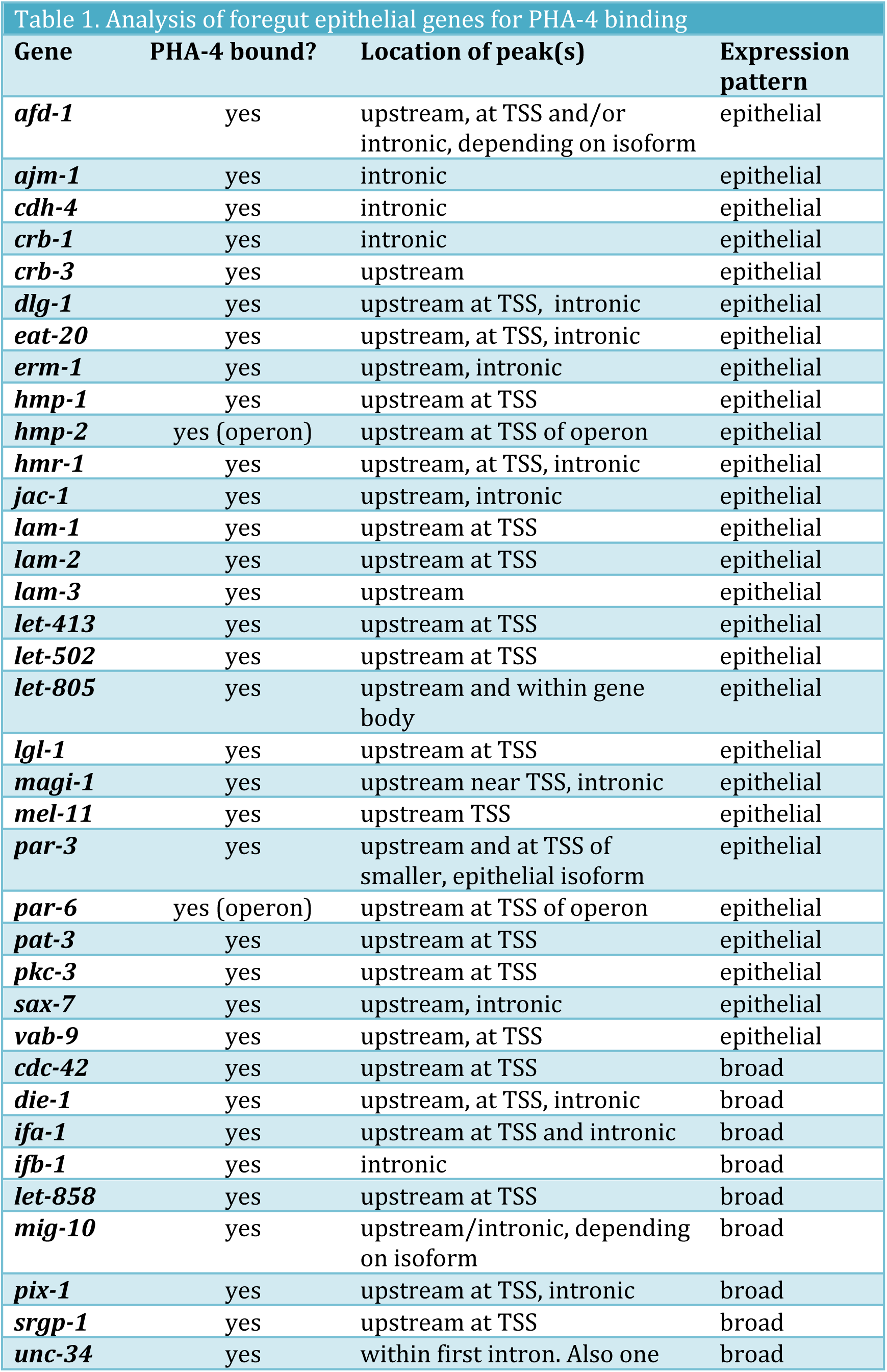
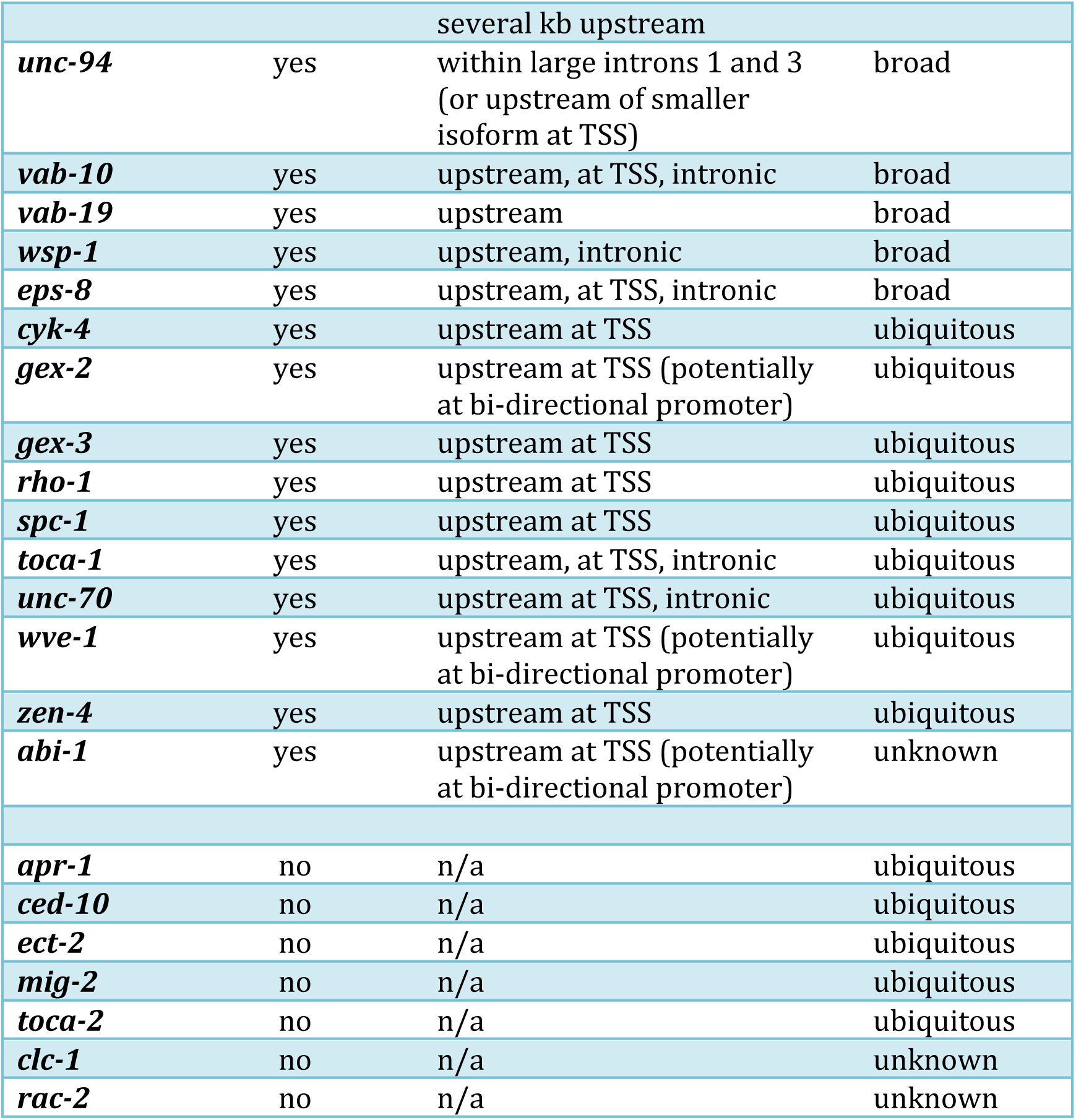
Analysis of foregut epithelial genes for PHA-4 binding

We examined other genes that have important functions in epithelial morphogenesis but a broad expression pattern within the embryo. 20/30 genes had clear PHA-4 peaks associated with cis-regulatory sequences, 3/30 had PHA-4 associated with the TSS of the polarity gene and the TSS of a neighboring gene on the opposite strand in a potentially bi-directional promoter, and 7/30 had no associated PHA-4 peak (Table 1). On the other hand, we rarely observed PHA-4 associated with genes that are expressed in other epithelia but not in foregut or Arcade Cells (only 1/12). For example, *ifb-2,* which codes for an apically-localized intermediate filament protein expressed in the midgut (Bossinger *et al.*, 2004), lacked PHA-4 bound to its regulatory sequences (Figure 5, Table S1). These findings suggest PHA-4 regulates a broad cohort of epithelial factors within the foregut and Arcade Cells.

### FoxA factors may regulate a conserved epithelial program

In mammals, FoxA factors promote epithelial organ development, and their loss often correlates with EMT of tumors. We asked whether FoxA factors target core epithelial factors in mammals as they do in *C. elegans*. We queried existing FoxA ChIP-seq datasets for mouse (Soccio *et al.*, 2011) and human cells (Motallebipour *et al.*, 2009; Hurtado *et al.*, 2011; Gosalia *et al.*, 2015; Wang *et al.*, 2015) to determine if FoxA factors bound to the regulatory elements of epithelial factors, examining 1000bp up and downstream of the gene body, as well as the gene body itself. We did not analyze enhancers, as they are often difficult to assign to a particular gene.

We examined homologues of the 50 epithelial-specific factors that were bound by PHA-4 in *C. elegans* (Table 1). 27/50 mouse epithelial homologs had an associated FoxA2 peak in primary liver cells (Table 2). Caco2 cells are derived from colorectal cancer but mimic normal polarized enterocytes when cultured to confluency (Fogh *et al.*, 1977; Alvarez-Hernandez *et al.*, 1991; Meunier *et al.*, 1995). Analysis of FOXA2 ChIP-Seq data revealed FOXA2 peaks near the human homologs of 33/50 *C. elegans* epithelial genes (Gosalia *et al.*, 2015). Intriguingly PARD3 and PARD3B - homologs of PAR-3, a master regulator of epithelial polarity - had 11 and 13 peaks throughout the gene body, mostly in introns, suggesting potential regulation by FoxA2. PARD3 and PARD3B were also bound by FoxA1 and FoxA2 in human embryonic stem cell-derived gut tube (GT) and foregut (FG) cells (Wang *et al.*, 2015). In GT and FG cells, respectively, FoxA1 bound 33/50 and 32/50 worm epithelial genes while FoxA2 bound 43/50 and 45/50 genes (Figure 6, Table 2). In addition to the *par-3* homologs listed above, mouse and human homologs of *dlg-1* and *erm-1* were also bound by FOXA1/2 in all datasets analyzed.

**Table 2.**
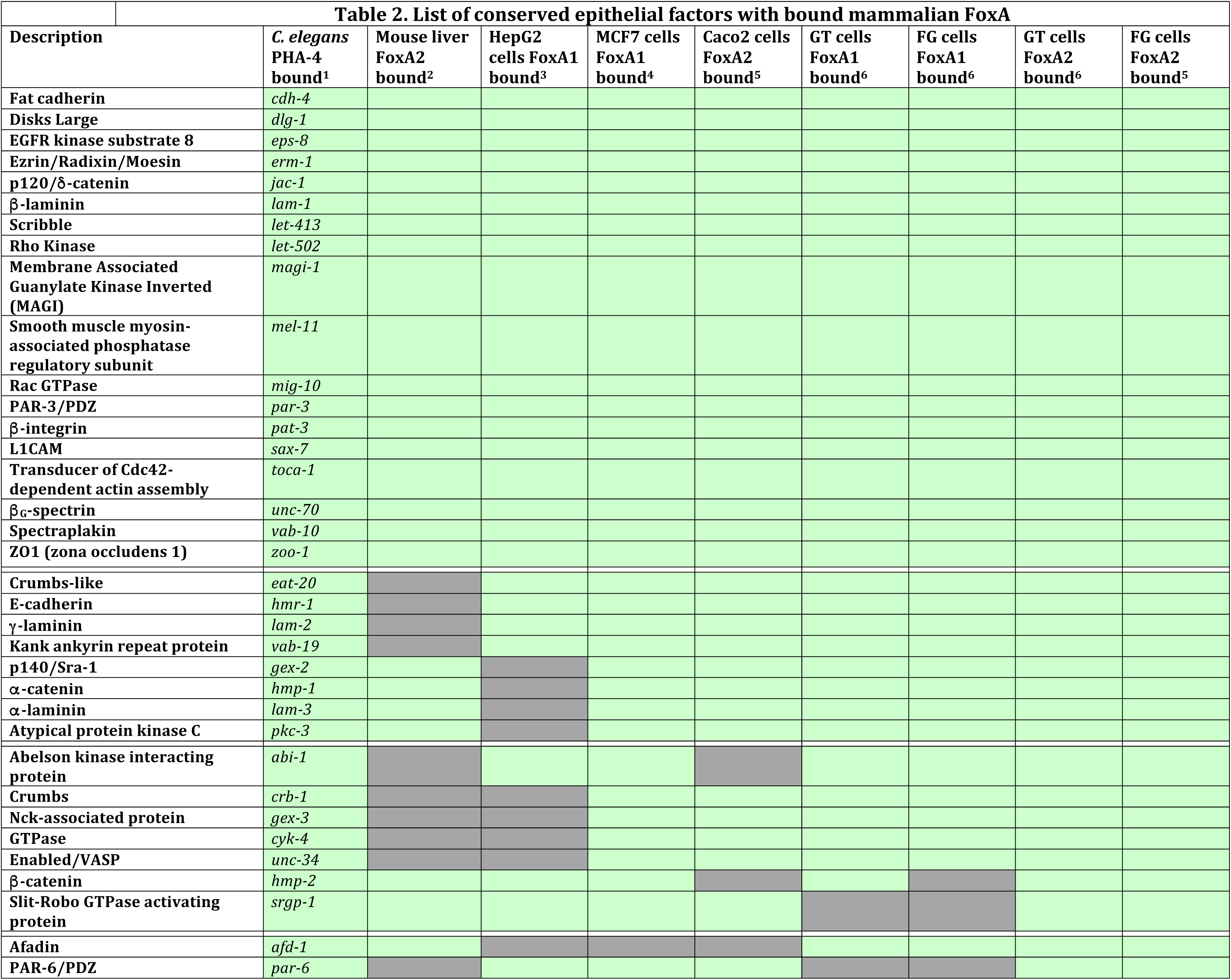
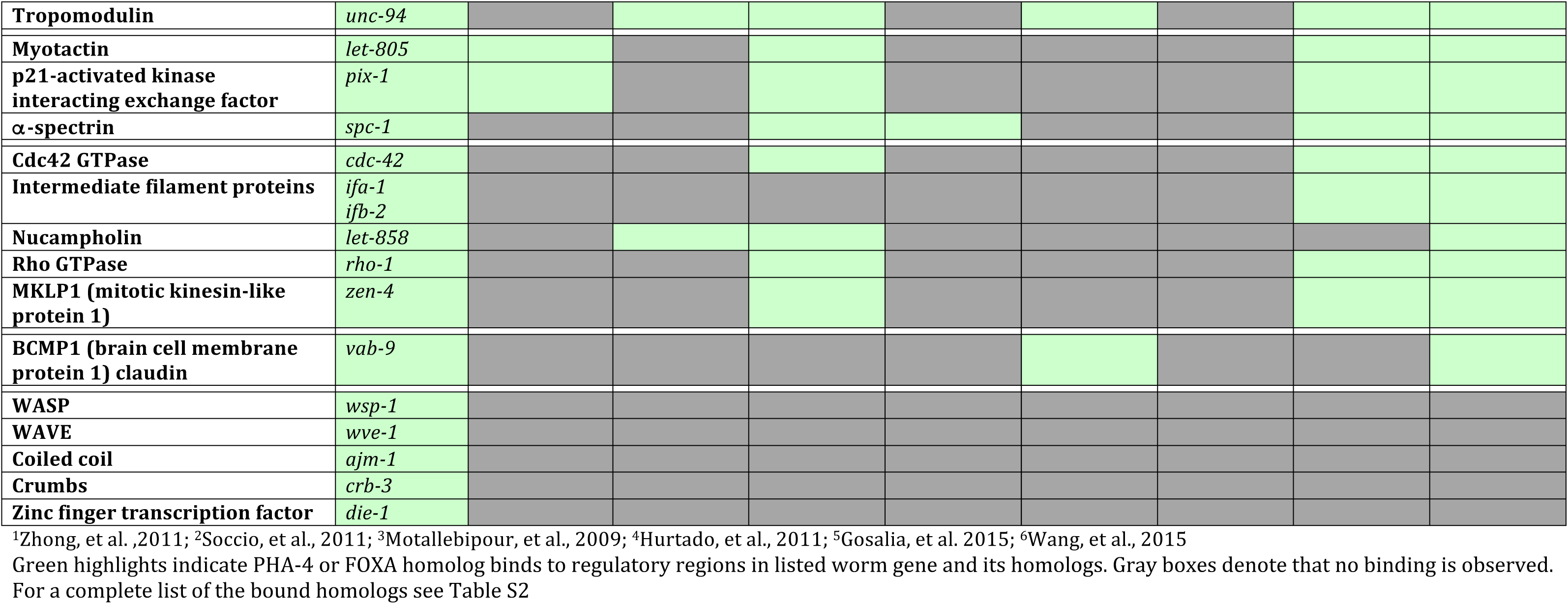
List of conserved epithelial factors with bound mammalian FoxA

**Figure 6.**
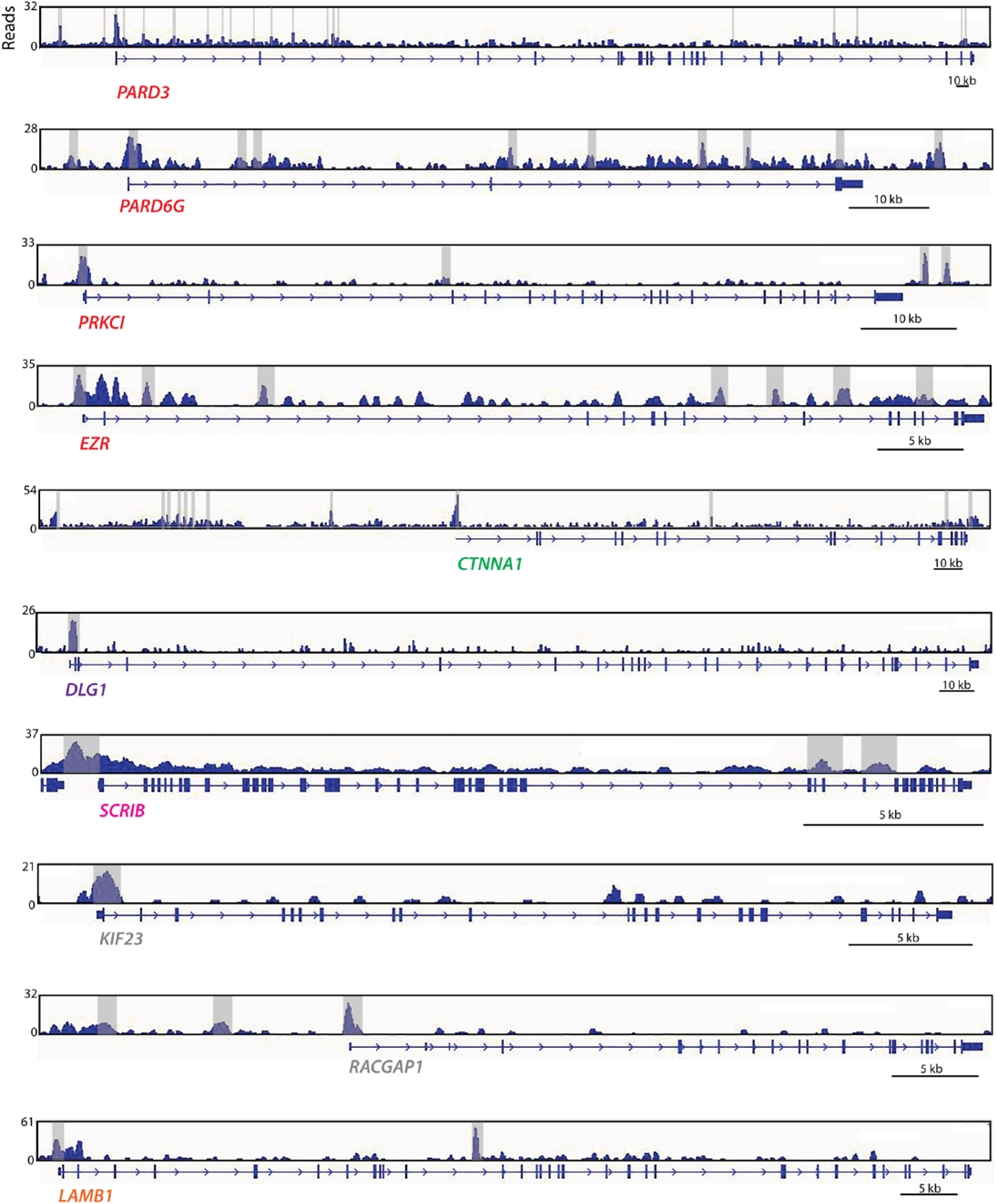
FOXA2 binds to promoter and intrans of mammalian epithelial genes. The genome browser panels display FOXA2 binding peaks for selected genes from the ChIP-Seq dataset generated by Wang et al (2011) from foregut-induced embrvonic stem cells. Significant FOXA2 peaks are highlighted in grav. Onlv a single representati ve gene model is shown, although several have alternatively spliced variants.

We also analyzed two datasets from human cancer cells derived from epithelial organs. HepG2 cells are derived from a hepatocellular cancer (Knowles *et al.*, 1980). FoxA1 binding was detected in 28/50 human homologs of nematode genes (Motallebipour *et al.*, 2009) (Table 2). Of these, 16 were shared between mouse liver and human liver cancer cells. MCF7 breast cancer cells were also examined, and we observed that FOXA1 bound 42/50 homologs of nematode epithelial genes (Hurtado *et al.*, 2011) (Table 2). In MCF7 cells, members of all three polarity groups were bound (*CRB1-2, PARD3/6, DLG1-3*), as were mediators of cell-cell (e.g. *CDH1*) and cell-matrix interactions (e.g. *ITGB1-8*).

In sum, of 50 gene families surveyed, FoxA1 or FoxA2 was observed in at least 7/8 datasets for 26 of the gene families (Table 2, Table S2). Included in this list are genes essential for proper epithelial polarity in mammalian cells (E-cadherin, β-integrin, Disks Large) as well as mediators of tight and adherens junction formation and function (Par3, ZO1, MAGI2, α-catenin, p120-catenin, Rho kinase). These results implicate mammalian FoxA for regulation of a common set of epithelial factors in many tissue types, and suggest that FoxA factors may function as master regulators of epithelial identity.

### ZEN-4 is required for protein accumulation in the arcade cell epithelium

Once transcription is activated throughout the digestive tract, the Arcade Cells delay accumulation of DLG-1 protein until polarity onset (Figure 2, comma stage). The ZEN-4 kinesin is essential for Arcade Cell polarity (Portereiko *et al.*, 2004), and we tested whether *zen-4* had a role in *dlg-1* accumulation. *zen-4* mutants had abundant *dlg-1* mRNA at the comma/1.25fold stages (when the Arcade Cells normally polarize), similar to wild-type siblings (Figure 7A, B). However, *zen-4* mutants lacked DLG-1 protein at the 1.25fold stage (compare Figure 7I to 7J). Maternally supplied *par-6* and *erm-1* gave similar results as *dlg-1*. Detectable *par-6* and *erm-1* RNA was observed in *zen-4* mutant arcade cells (Figure 7D, F) but there was no detectable protein (Figure 7 H, L). Protein was also missing for previously assayed apical (PKC-3, PAR-3) and junctional (AJM-1, HMR-1) markers (Portereiko *et al.*, 2004). We conclude that *zen-4* is critical to accumulate the polarity proteins required to form the Arcade Cell epithelium.

**Figure 7.**
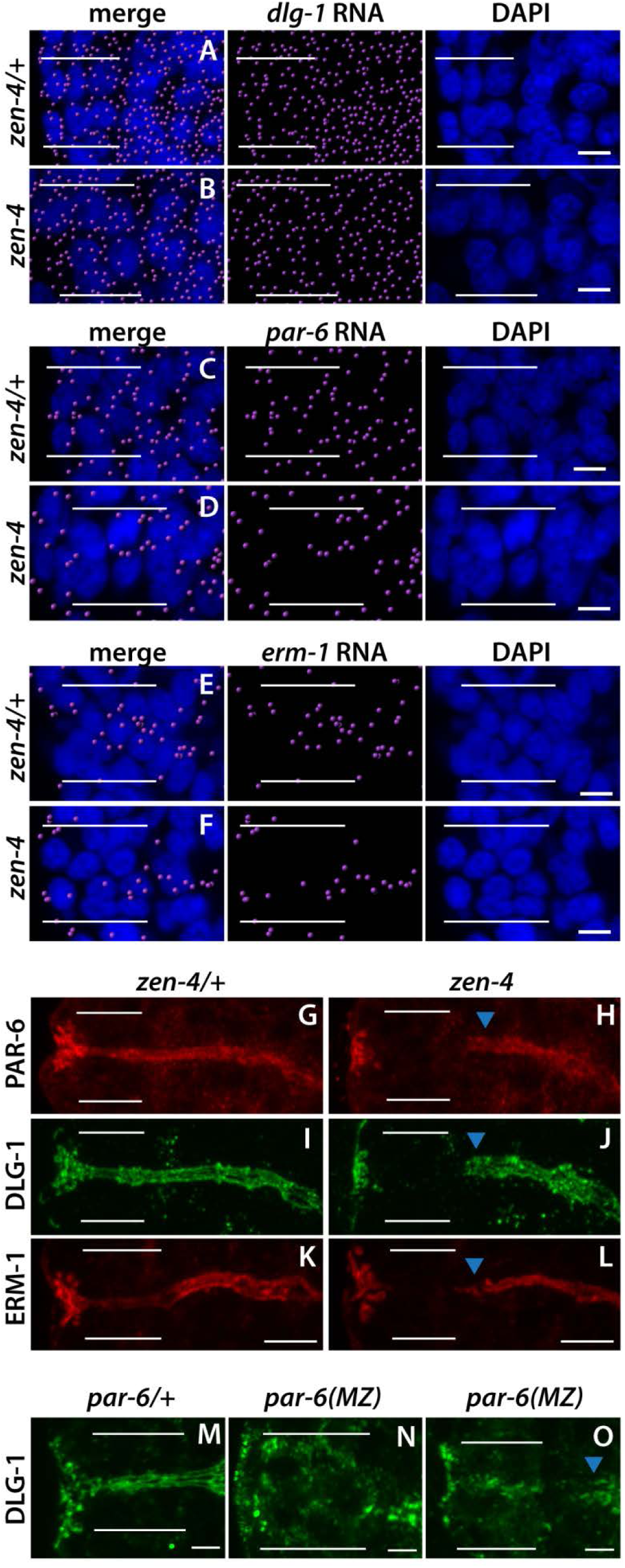
zen-4 and par-6 are required for Arcade Cell Polarization. A-F. mRNA detected by smFISH pseudo-colored magenta, DAPI is blue. Epithelial factor mRNAs (dlg-1, par-6, erm-1) are present in both zen-4/+ (A, C, E) and zen-4 mutant (B, D, F) Arcade Cells, which are highlighted with white bars. G-L. PAR-6 (G), DLG-1 (I), and ERM-1 (K) are expressed and localized to the apical/junctional domain in zen-4/+ Arcade Cells (white bars). These proteins are not detected in zen-4 mutant Arcade Cells (H, J, L), although they are expressed and localized in the neighboring foregut epithelium (blue arrowhead). M-O. DLG-1 (green) is expressed and properly localized in the arcade cells (white bars) of par-6/+ (M) embryos. In 407% (14/35) of par-6 mutant embryos DLG-1 is detected but fails to localize properly within the arcade cells (N). In the other 60% (21/35) DLG-1 is properly localized but fails to form continuous junctions, similar to the neighboring foregut (blue arrowhead) (O). All images are maximum intensity projections. Anterior is to the left. Scale bars are 2 µm (M-O) or 5 µm (G-L).

The effect of *zen-4* on accumulation of polarity proteins rather than localization suggested ZEN-4 would function early, prior to polarity onset. To test this idea, we performed temperature-shift experiments to pinpoint when *zen-4* activity was required (Figure 8). We used a temperature-sensitive allele of *zen-4, or153ts* (Severson *et al.*, 2000), which abrogates binding to its obligate partner in cytokinesis, *cyk-4* when shifted to the restrictive temperature (Pavicic-Kaltenbrunner *et al.*, 2007). We inactivated *zen-4* during three time periods: close to the time of Arcade Cell birth (early); during Arcade Cell polarization (late); and between the early and late time points (middle). Embryos were shifted to restrictive temperature for a 70-minute window followed by incubation at permissive temperature to the 1.5 fold stage, when the Arcade Cells have formed an epithelium in the wild type. We confirmed rapid inactivation of *zen-4* at these late stages by testing for cytokinesis defects (Figure S5), which is a canonical phenotype of *or153ts* (Severson *et al.*, 2000).

**Figure 8.**
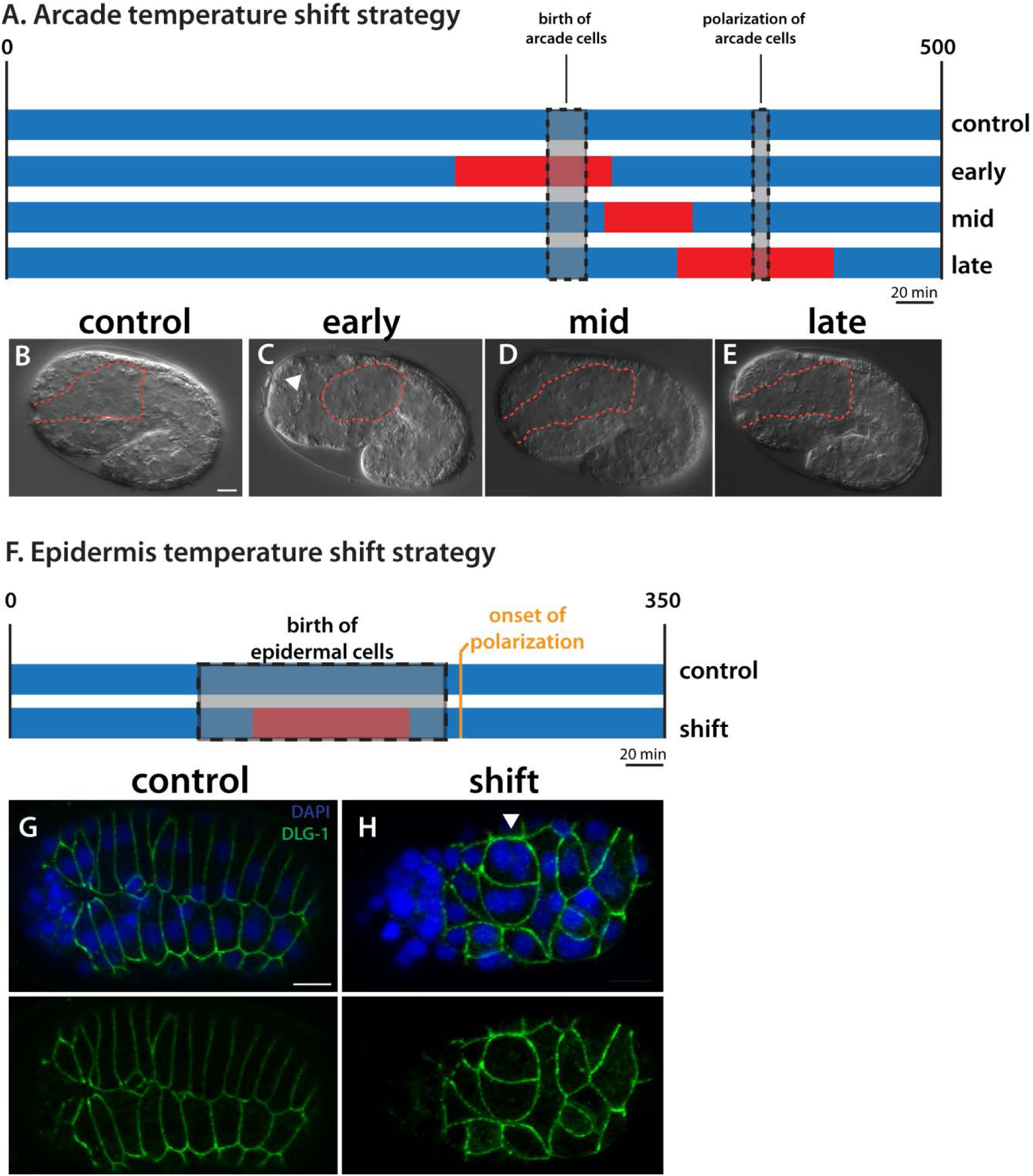
*zen-4* temperature-sensitive experiments show that ZEN-4 activity is required at Arcade Cell birth. A. Temperature-shift strategy. Blue represents permissive temperature (15°C) and red denotes restrictive temperature (26°C). The control sample was ubated at the permissive temperature only. The early shift was started ~50 minutes prior to Arcade Cell birth (grey box), while late shift was started ~50 minutes prior to Arcade Cell polarization (grey box). The mid shift occurred in between the early and e shifts. B-E. Representative images of embryos after the above incubations. The foregut-associated basement membrane (BM) is hlighted by a red dotted line. The control, mid and late embryos all have a polarized arcade cell epithelium, as evidenced by ving attached foreguts. Note the BM surrounds an elongated foregut (red). The early shift results in an unattached foregut cause the arcade cells (arrowhead) have failed to polarize. Note the BM surrounds the whole foregut primordium in the center of embryo (red). F. Early temperature-shift strategy to inactivate ZEN-4 during epidermal cell division. Epidermal cells are born er a longer period of time than the Arcade Cells (grey box), and this period encompasses the time at the restrictive temperature d). The embryos were shifted to permissive temperature prior to the onset of epidermal polarization. G, H. Representative ximum intensity projections through the epidermis of embryos at the end of the shift. In the control embryo (G), DLG-1 (green) is nd at the cell junctions of epidermal cells that have formed two rows on the dorsal surface. These cells appear mono-nucleate. In shifted embryo (H), the epidermal cells still generate and localize DLG-1 properly, even though many cells are obviously multi-nuleate (arrowhead) due to cytokinesis defects. Anterior is to the left. Scale bars are 5 µm.

The majority of embryos shifted to non-permissive temperature at the time of Arcade Cell birth had defects in polarization (84%; 27/32). On the other hand, almost all embryos shifted during the middle and late periods appeared wild-type (35/37 and 55/56 for the middle and late windows, respectively). These data suggest that ZEN-4 is required around the time of Arcade Cell birth, rather than at the time of polarization, consistent with the idea that ZEN-4 acts early to produce polarity proteins, which are later assembled into apical and basolateral domains. As a control, inactivation of *zen-4(ts)* at the birth of the epidermal cells did not disrupt polarity in the epidermis, indicating *zen-4* is required specifically in the Arcade Cells and that blocked cytokinesis is not sufficient to disrupt epithelial polarity (Figure 8).

### Structure-Function Analysis of ZEN-4 reveals a requirement for binding to CYK-4/GAP

ZEN-4/MKLP relies on four protein domains to fulfill its roles in cytokinesis and neurite outgrowth (White and Glotzer, 2012). We performed structure-function analysis to gain insight into which domains were important for Arcade Cell polarization (Figure 9A). We assayed both rescue of the *zen-4(px47)* mutant phenotype (see Methods) as well as localization of a GFP-tagged reporter. To monitor *px47* rescue, we examined progeny from heterozygous *zen-4(px47)* mothers. Normally 25% offspring are *zen-4(px47)* homozygotes, and of these, half or more arrest with unpolarized Arcade Cells (Table 4, Table S3). A wild-type ZEN-4::GFP transgene rescued homozygotes, producing 0% embryonic lethality from heterozygous mothers (Figure 9A and Table S3). These animals displayed a dynamic localization pattern for ZEN-4::GFP that mimicked endogenous ZEN-4 and its homologs (Figure 9B; (Powers *et al.*, 1998; Raich *et al.*, 1998; Deavours and Walker, 1999; Chen *et al.*, 2002; Minestrini *et al.*, 2002, 2003; Verbrugghe and White, 2007)). ZEN-4::GFP was nuclear in interphase cells, localized to the ingressing furrow and cortex during mitosis, and remained at the division remnant after abcission.

**Figure 9.**
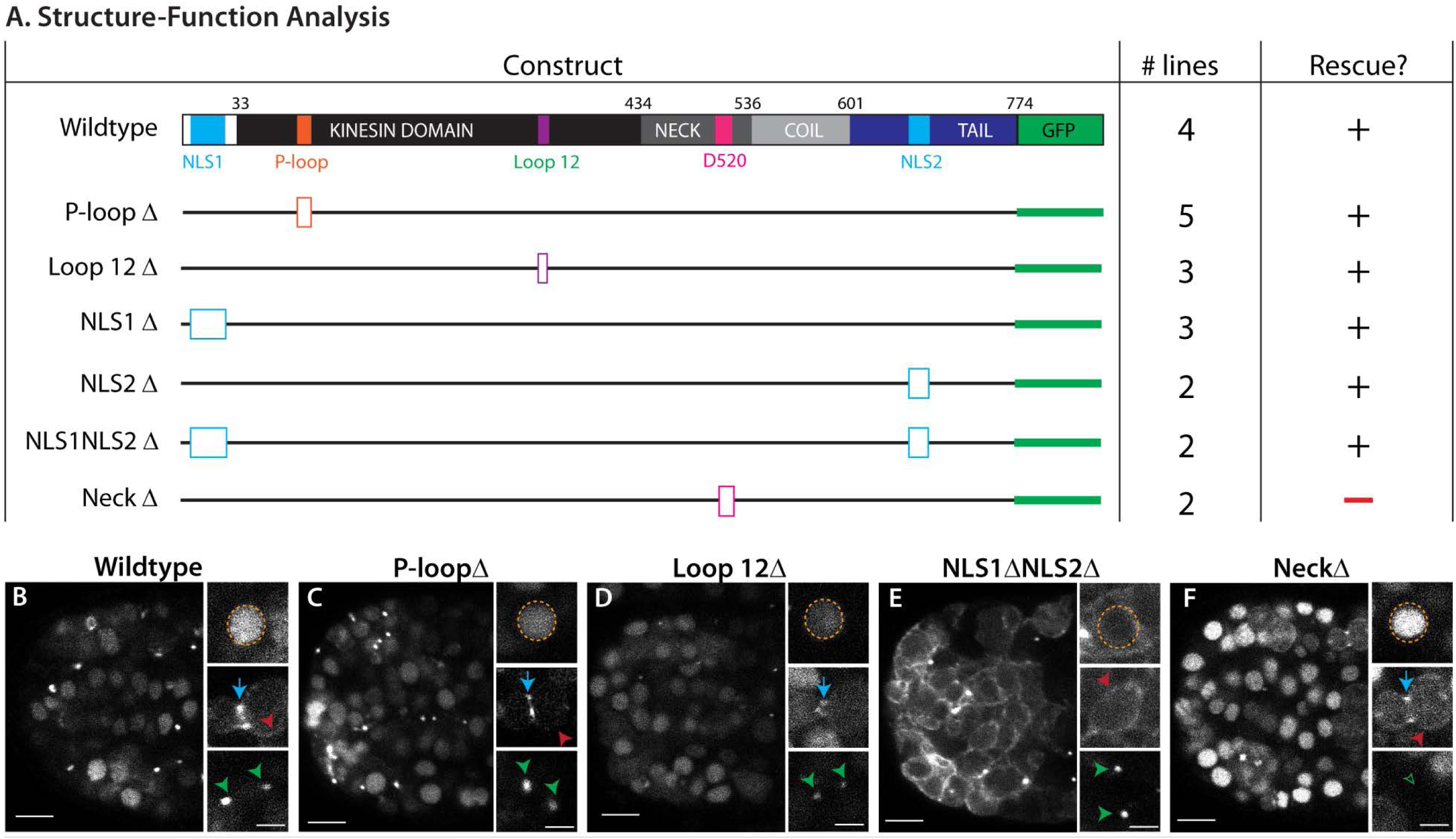
ZEN-4 structure-function analysis reveals a requirement for residues that bind CYK-4/MgcRacGAP. A. The domain structure of wild-type ZEN-4 is shown at the top. There are two nuclear localization sequences (NLS, light blue), one at the N-terminus and one in the tail domain (dark blue). The Kinesin domain (black) contains the ATP-binding pocket known as the P-loop (orange), and loop 12 (purple), which is important for binding to microtubules. The neck domain (dark gray) contains residues important for binding to CYK-4/MgcRacGAP, including D520. There is also a coiled-coil domain (light gray). GFP is linked to the C-terminus and is shown in green. To test the function of each domain in polarity we generated small deletions (open boxes) to remove key residues for each domain. The number of transgenic lines expressing each construct is listed (# lines). A plus sign designates if the construct could rescue the *zen-4(px47)* Arcade Cell polarization defect, while a minus sign denotes a lack of rescue. See Tables 3 and S3 for numbers). B-F. Live-imaging of ZEN-4::GFP at the 16E stage, focused on the foregut/arcade cells. B. Wildtype ZEN-4::GFP is mostly nuclear at this stage (orange circle), but is also visible in the furrow (blue arrow) and cortex (red arrowhead) of dividing cells, and at the division remnant (green arrowhead). C, D. The kinesin mutants (P-loopΔ, Loop 12Δ) have similar localization to wild-type. E. The double NLS deletion (NLS1ΔNLS2Δ) shows constitutively cortical localization of ZEN-4, is still apparent in division remnants, and is much less enriched in the nucleus, although faint signal is still apparent. F. The NeckΔ is still apparent in the nucleus and at the furrow/cortex, but localization to the division remnant is lost (open green arrowhead). Anterior is to the left. Scale bars are 5µm (overview) and 2µm (closeups).

**Table 3.** ZEN-4 structure-function analysis

| **Construct** | **Array** | **transgene negative**<br>**% unpolarized (n)** | **transgene positive**<br>**% unpolarized (n)** |
| --- | --- | --- | --- |
| **Wildtype** | *pxEx390* | 15% (17/115) | 0.7% (1/150) |
|  | *pxEx392* | 15% (8/54) | 0% (0/92) |
|  | *pxEx393* | 12% (8/67) | 0% (0/136) |
|  | *pxEx396* | 16% (18/115) | 0% (0/128) |
| **P-loopΔ** | *pxEx395* | 20% (12/60) | 0% (0/52) |
|  | *pxEx399* | 12% (22/181) | 2% (5/246) |
|  | *pxEx402* | 15% (16/107) | 0% (0/89) |
|  | *pxEx486* | 17% (14/83) | 1% (1/82) |
|  | *pxEx488* | 13% (4/29) | 1% (1/72) |
| **Loop12Δ** | *pxEx422* | 18% (25/136) | 1.5% (2/129) |
|  | *pxEx424* | 17% (30/181) | 0% (0/363) |
|  | *pxEx489* | 14% (7/51) | 0% (0/157) |
| **NLS1Δ** | *pxEx397* | 18% (8/44) | 2% (2/82) |
|  | *pxEx423* | 16% (16/101) | 0% (0/67) |
|  | *pxEx425* | 16% (6/41) | 5% (2/43) |
| **NLS2Δ** | *pxEx398* | 18% (58/324) | 0.6% (2/327) |
|  | *pxEx420* | 19% (13/67) | 0% (0/94) |
| **NLS1/2Δ** | *pxEx455* | 16% (15/91) | 1.5% (2/134) |
|  | *pxEx456* | 11% (12/110) | 0% (01/132) |
| **NeckΔ** | *pxEx391* | 17% (32/189) | 20% (44/222) |
|  | *pxEx444* | 16% (39/119) | 19% (47/249) |

**Table 4.** Tethered dimer analysis

| Table 4. Tethered dimer analysis |  |  |  |
| --- | --- | --- | --- |
| Construct | Array | transgene negative<br>% unpolarized (n) | transgene positive<br>% unpolarized (n) |
| ZEN-4::CYK-4 | *pxEx518* | 10% (9/96) | 0.6% (1/170) |
|  | *pxEx519* | 16% (13/80) | 3.6% (5/137) |
|  | *pxEx541* | 13% (22/171) | 0.5% (1/206) |
| ZEN-4(NeckΔ)<br>::CYK-4 | *pxEx516* | 11% (12/109) | 0.1% (1/74) |
|  | *pxEx517* | 12% (46/398) | 0.1% (3/285) |
| ZEN-4::AIR-2 | *pxEx542* | 15% (34/228) | 0.7% (1/145) |
|  | *pxEx543* | 18% (6/33) | 0% (0/135) |
|  | *pxEx544* | 22% (14/63) | 4% (3/82) |
| ZEN-4(NeckΔ)<br>::AIR-2 | *pxEx523* | 20% (26/128) | 13% (25/194) |
|  | *pxEx524* | 12% (17/142) | 13% (20/155) |
|  | *pxEx556* | 22% (22/101) | 15% (25/172) |

ZEN-4(NeckΔ)::GFP lacks 5 residues around residue D520 within the neck domain. This construct was unable to rescue *zen-4(px47)*: 0/2 transgenic ZEN-4(NeckΔ)::GFP lines rescued *zen-4(px47)* mutants, like the no-transgene control (Figure 9B, Table 4, Table S3). The neck domain serves two purposes. It is essential to bind CYK-4/MgcRacGAP, a GTPase activating protein that controls ZEN-4 kinesin activity (Mishima *et al.*, 2002; Pavicic-Kaltenbrunner *et al.*, 2007; White *et al.*, 2013). It is also important to localize ZEN-4 to the cell cortex and division remnant. To determine if ZEN-4 requires CYK-4 activity or division remnant localization, we tethered ZEN-4(NeckΔ) to CYK-4 using a flexible linker (see Methods) (Neuhold and Wold, 1993; Castanon *et al.*, 2001). As a positive control, we linked CYK-4 to wild-type ZEN-4. As shown in Figure 10A and Table S3, both wild-type and mutant fusions were sufficient for rescue, indicating that association with CYK-4 is critical for ZEN-4 function and disrupted by the NeckΔ mutation.

**Figure 10.**
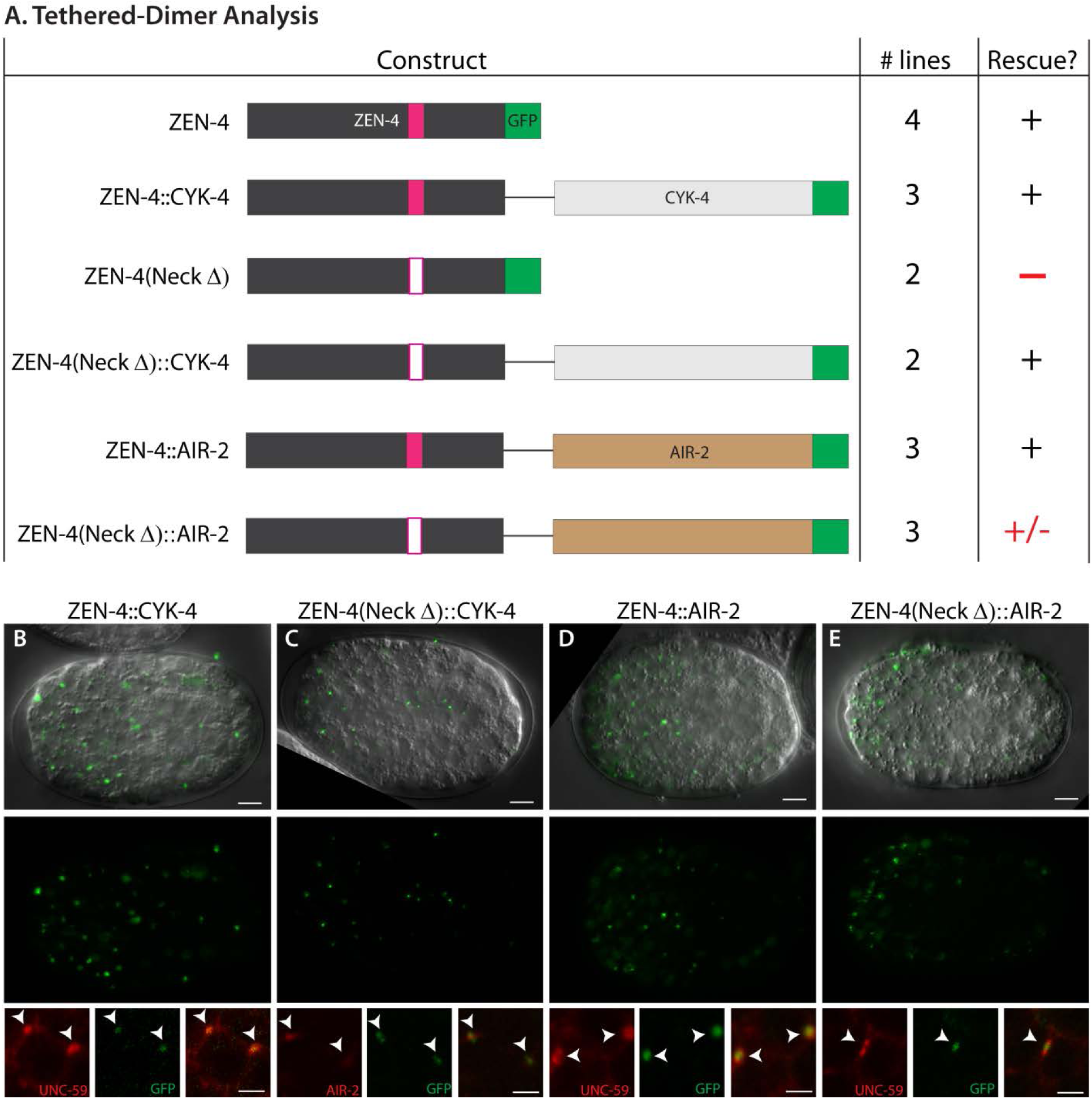
ZEN-4 tethered dimer analysis demonstrates the importance of the ZEN-4/CYK-4 interaction. A. Wildtype ZEN-4 or the ZEN-4(NeckΔ) were tethered to either GFP-tagged CYK-4 or AIR-2 (AuroraB kinase) via a flexible linker (black line). The number of lines and ability to rescue the *zen-4(px47)* arcade cell polarization defect are shown (See Tables 4 and S4 for numbers). Untethered ZEN-4 and ZEN-4(NeckΔ) from Figure 9 are repeated here for ease of reference. B-E. Live-imaging of GFP-tagged ZEN-4 tether constructs at the 16E stage. The majority of signal is found at division remnants (white arrowheads), identified by division remnant markers UNC-59/Septin (Nguyen *et al.*, 2000) or AIR-2 (Schumacher *et al.*, 1998). UNC-59 is localized to the cell cortex but becomes enriched at the division remnant following furrow ingression. Anterior is to the left. Scale bars are 5µm (whole embryo) and 2µm (closeups).

The tethered dimer localized ZEN-4::CYK-4 to the division remnant. To determine if localization was sufficient for rescue, we tethered ZEN-4 to the aurora kinase AIR-2/AuroraB, which is enriched at the division remnant partially overlapping with CYK-4 (Schumacher *et al.*, 1998) and is required to localize ZEN-4 normally (Severson 2000). A wild-type ZEN-4::AIR-2 fusion served as a control. AIR-2 was only weakly able to rescue ZEN-4(NeckΔ) despite complete localization to the division remnant (Figure 10, Table 5). Embryos bearing ZEN-4(NeckΔ)::AIR-2 had 18% lethality (range 17-19%), compared to ~23% lethality for no-transgene control embryos (range 17-27%). The wild-type ZEN-4::AIR-2 control was fully functional (0-4% lethality for array+ embryos), demonstrating that AIR-2 did not interfere with ZEN-4 function. We conclude that accumulation at the division remnant is not sufficient for rescue, and that recruitment of CYK-4 is important independent of its effects on localization.

CYK-4 can modulate actin dynamics by way of its GTPase activating domain (Canman *et al.*, 2008; Loria *et al.*, 2012; Zhang and Glotzer, 2015). It also activates the ATPase and kinesin activities of ZEN-4 on microtubules (Mishima *et al.*, 2002; White *et al.*, 2013). To determine whether microtubules were critical for ZEN-4 function in polarity, we mutated the P loop and loop 12 within the kinesin domain. The P-loop contains the ATPase activity of the kinesin (Saraste *et al.*, 1990; Nakata and Hirokawa, 1995), and mutations in this domain in ZEN-4 homologs result in a motor-dead protein that can still bind microtubules (Chen *et al.*, 2002; Matuliene and Kuriyama, 2002). Loop 12 of kinesins is important for binding to microtubules (Woehlke *et al.*, 1997; Raich *et al.*, 1998). These mutations had a mild defect on ZEN-4 localization and did not impact rescue (Figure 9 C, D). Thus, we have no evidence that microtubule activities are important for ZEN-4 during polarity. Finally, we note that ZEN-4 normally localizes to nuclei during interphase. Disruption of two predicted nuclear localization signals (NLSs) strongly reduced nuclear accumulation but had no observable effect on rescue (Figure 9E, Table 4). Similarly, mutation of either NLS1 or NLS2 alone did not affect ZEN-4 activity (Figure 9A, Table 4). These data suggest that neither the kinesin function nor nuclear localization are necessary for polarity, although we cannot rule out that residual ZEN-4 in the nucleus or movement on microtubules is sufficient for rescue. In summary, ZEN-4 relies on its interaction with CYK-4 to induce polarity. The ATPase and kinesin activities appear unimportant, suggesting that regulation of actin may be central for controlling protein accumulation of polarity proteins. In agreement, actin is severely disrupted in *zen-4(px47)* mutant Arcade Cells (Portereiko *et al.*, 2004).

### PAR-6 is required to position DLG-1 within nascent Arcade Cell epithelia

The data suggest that PHA-4 and ZEN-4 control RNA and protein production of polarity factors in the Arcade Cells. Once produced, standard polarity pathways likely organize the apical, junctional and basolateral domains within the Arcade Cells (Rodriguez-Boulan and Macara, 2014). We tested this idea by examining *par-6* mutants. Because *par-6* is important for the first embryonic cell divisions, we removed PAR-6 protein at the gastrula stage, using the elegant ZF1 system of Nance and colleagues (*par-6m+z-*) (Nance *et al.*, 2003). *par-6* mutants produced DLG-1 protein but often failed to localize it properly to the junctional domains (Figure 7). Instead, we detected DLG-1 in the cytoplasm of *par-6(m+z-)* mutants (Figure 7N). These data reveal that temporal regulation of the Arcade Cell epithelium depends on three levels of tissue-specific control: first at the transcriptional level, second at the level of protein expression and third at the level of protein localization to nascent adherens junctions.

## DISCUSSION

The last three decades have seen major advancements in our understanding of cell polarization during mesenchymal-to-epithelial transition (MET) (Chaffer *et al.*, 2007), however little is known about coordination between distinct epithelia to produce the mature body plan. Our data show that *C. elegans* epithelia polarize just prior to the morphogenetic event in which that epithelium is involved (e.g. epiboly, torus formation). In particular, the Arcade Cells delay polarization until after the epidermis and gut epithelia have matured sufficiently to withstand the pulling forces of embryo elongation. We have discovered four regulators that control this delayed timing: PHA-4/FoxA initiates expression of polarity genes within the Arcade Cells, ZEN-4/MKLP1 and CYK-4/MgcRacGAP dictate the onset of protein accumulation, and PAR-6 controls subcellular localization within the nascent epithelium.

### PHA-4/FoxA regulates a complete epithelial program

The FoxA pioneer transcription factor PHA-4 promotes the onset of *dlg-1* expression within the foregut and Arcade Cells, but not other cell types. In addition to *dlg-1*, the majority of apical, junctional, and basolateral genes had an associated PHA-4 ChIP peak, most either associated with the TSS or within the first intron, which are hallmarks of bona fide PHA-4 target genes (Gaudet and Mango, 2002; Gaudet *et al.*, 2004). All of these genes are also expressed in other epithelia (or other cell types). We confirmed that expression for two of these factors (PAR-3, ERM-1) was selectively reduced in the foregut and the Arcades. This observation rules out the possibility of a single epithelial program regulated by a common set of transcription factors in all epithelial organs, and suggests that the mechanism of timing is controlled at the tissue and organ level by tissue/organ-specific transcription factors such as PHA-4.

The *P*_*4.9*_*dlg-1* construct was able to recapitulate the endogenous expression of DLG-1 protein in the foregut/arcade cells and the epidermis (Figure 4). A recent study of the *C. elegans* transcriptional start site and enhancer landscape identified a putative enhancer for *dlg-1* about 6kb upstream of the ATG start (Chen *et al.*, 2013). Our analysis revealed that this enhancer was not necessary for normal *dlg-1* expression in the foregut/arcade cells, at least in the context of a complex multi-copy array. It is possible that this region functions as a shadow enhancer that promotes robust *dlg-1* expression but is not essential (Hong *et al.*, 2008; Perry *et al.*, 2010). In addition, the *P*_*4.9*_*dlg-1* construct carried a heterologous 3’UTR, suggesting that the endogenous 3’UTR is not necessary for proper expression in the foregut and Arcade Cells. Perhaps regulation occurs through the 5’UTR, as has been suggested for some post-transcriptional events in *C. elegans* (Lee and Schedl, 2004; Jungkamp *et al.*, 2011; Vora *et al.*, 2013). We note that in the midgut, some cells expressed *P*_*4.9*_*dlg-1* RNA one cell-cycle earlier (data not shown), suggesting additional cis-regulatory sites for this tissue.

### FOXA regulation of a general epithelial program may be conserved

We extended our analysis to two mammalian homologs of PHA-4, FOXA1 and FOXA2. FOXA factors are necessary for the proper development of many epithelial organs, mostly of endodermal origin (e.g. liver, pancreas, intestine, lung). Conversely, epithelial tumors often lose FoxA expression during EMT (Bersaas 2016; Tang 2011 Cell Research, Song 2010). These studies have suggested that FoxA acts by suppressing the transcription factor Slug, a known regulator of EMT, other transcription factors and adhesion signaling pathways (Song *et al.*, 2010; Tang *et al.*, 2011; Wang *et al.*, 2014). Our survey revealed FOXA1/FOXA2 binding to the promoter and introns of mammalian homologs of the 50 epithelial-specific worm genes with PHA-4 binding (Figure 5, Table 2, Table S1), suggesting that FOXA1/A2 may promote a general epithelial program in mammals as in worms.

### ZEN-4 and PAR-6 control polarity onset

Our analysis revealed a delay between the appearance of *dlg-1* transcripts and the accumulation of protein within the Arcade Cells. ZEN-4 was critical for this regulation, as transcripts were abundant in *zen-4* mutant Arcade Cells, but protein was absent for 3/3 genes tested. We note that absence of protein was not due to mis-localization within cells because *par-6* mutants displayed bountiful mis-localized DLG-1 (this work) and ERM-1 (Von Stetina and Mango, 2015). Instead, ZEN-4 likely regulates the translation or stability of DLG-1 and other polarity proteins. This role suggests that ZEN-4 acts upstream of PAR-6, which is in agreement with the absence of PAR-6 in *zen-4* mutants and the temperature-sensitive period of *zen-4(ts)* mutants. ZEN-4 is a kinesin and possesses multiple structural domains. Kinesin activity and nuclear localization appeared to be dispensable for ZEN-4 function, but binding to CYK-4/GAP was essential. These data suggest that regulation of actin rather than microtubules may be the critical parameter for polarization. Consistent with this idea, actin, but not tubulin, is dramatically disrupted in *zen-4* mutant Arcade Cells (Portereiko *et al.*, 2004, data not shown). Previous studies have revealed that ZEN-4 and CYK-4 can modulate actin dynamics by regulating RhoA and Rac activity (Jenkins *et al.*, 2006; Canman *et al.*, 2008; Loria *et al.*, 2012; Zhang and Glotzer, 2015).

Tethering of mutant ZEN-4 to the division remnant gave either full (CYK-4) or partial (AIR-2) rescue. This result raises the intriguing possibility that localization to the remnant may contribute towards ZEN-4 function. The division remnant has been implicated in other polarizing events such as the positioning of dendrites in neuronal cells or establishing the dorsoventral axis in *C. elegans* (Pollarolo *et al.*, 2011; Singh and Pohl, 2014; Dionne *et al.*, 2015). Marking a site between dividing cells has been studied most extensively in yeast, where the bud scar serves as a marker for the next division (reviewed in Casamayor and Snyder, 2002; Slaughter *et al.*, 2009; Bi *et al.*, 2012).

The data presented here offer a possible explanation for the selectivity of *zen-4* for the Arcade Cells. Only the Arcade Cell epithelium is disrupted in *zen-4* mutants; other epithelia polarize normally (Portereiko *et al.*, 2004; Hardin *et al.*, 2008). Wild-type Arcade Cells exhibited a ~100-minute delay between the onset of *dlg-1* RNA and protein, whereas other epithelia displayed no delay or a much shorter delay. An appealing hypothesis is that the Arcade Cells depend on an extra layer of post-transcriptional regulation, mediated by ZEN-4 and CYK-4, compared to other embryonic epithelia. This extra layer of regulation may be critical to ensure polarization occurs late in embryogenesis.

Homologues of ZEN-4 and CYK-4 have been implicated in polarity at later stages of epithelium formation. These factors modulate RhoA and Rac1 in epithelial junctions of vertebrate cells (Ratheesh *et al.*, 2012; Guillemot *et al.*, 2014; Breznau *et al.*, 2015). This activity may be missing or redundant in *C. elegans*, because conditional inactivation of *zen-4* did not reveal a requirement at mid or late time points. We note that the vertebrate studies did not design their experiments to reveal an early role in polarity protein accumulation, as described here, so it remains to be seen whether ZEN-4 function is conserved.

Once DLG-1 and other polarity proteins are produced, they are rapidly localized within the cell. We found that *par-6* was necessary for localization of DLG-1 within the Arcade Cells in at least 50% of tested embryos. These data extend our previous observations that in half of the tested embryos, several markers of epithelial polarity (actin, HMR-1/cadherin, ERM-1, AJM-1) failed to localize properly within the Arcade Cells (Von Stetina and Mango, 2015). In the other half, protein is localized properly (like DLG-1) but fails to form continuous junctions between other Arcade Cells and between tissues (i.e. foregut and arcades). In other epithelia, this failure to mature junctions is the most severe *par-6* phenotype observed but cell polarity is intact (Totong *et al.*, 2007; Von Stetina and Mango, 2015). What accounts for the difference? One possibility is that distinct tissues are differentially dependent on specific proteins. For example, PAR-6 is more important in the epidermis than PAR-3, while *par-3* mutants have a more severe phenotype in the midgut than *par-6* mutants (Achilleos *et al.*, 2010). By this logic, PAR-6 may be a more important player for polarization in Arcade Cells compared to other tissues. An alternative possibility is that residual PAR-6 may provide polarizing activities. The M/Z system removes wild-type protein expressed from maternally-deposited transcripts, but the system takes time to complete. Perhaps residual PAR-6 is present and active when other epithelia are formed, but gone by the time the Arcade Cells undergo polarization. Consistent with this idea, embryos with partial Arcade Cell polarity always exhibited some PAR-6 protein, whereas those that did not lacked detectable PAR-6 completely (Von Stetina and Mango, 2015).

## Materials and methods

### Strains and maintenance

Nematodes were grown at 20°C using standard conditions (Brenner, 1974), unless otherwise stated. Some strains were provided by the CGC. Strains used: N2 (SM1880); SM469 [*pxIs6* (*pha-4*::GFP::Histone)] (Portereiko and Mango, 2001); SM1052 [*zen-4 (px47) dpy-20 (e1282)/bli-6(sc16) unc-24(e138)* IV] (Portereiko *et al.*, 2004); FT250 [*unc-119(ed3)* III; *xnIs96* (HMR-1::GFP)] (Achilleos *et al.*, 2010); FZ223 [*mcIs204* (*P_7_dlg-1*::DLG-1::GFP; pRF4)] (McMahon *et al.*, 2001); JJ1743 [*par-6(tm1425)/hIn1[unc-54(h1040)]* I*; him-8(e1489)* IV] (Totong *et al.*, 2007); FT36 [*unc-101(m1) par-6(zu170)*I; *zuIs43* (*pie-1*::GFP::PAR-6::ZF1 + *unc-119*(+))] (Totong *et al.*, 2007); EU554 [*zen-4(or153*) IV] (Severson *et al.*, 2000). Strains generated for this study can be found in Table S5.

### Temperature-shift Experiments

*zen-4(or153ts)* was maintained at 15°C. To perform arcade cell temperature-shifts on staged embryos, gravid adults were transferred to a pre-cooled (4°C) dissection chamber with 50ul ice-cold M9 Buffer (22 mM KH_2_PO_4_, 22 mM Na_2_HPO_4_, 85 mM NaCl, 1 mM MgSO_4_). Embryos were released by dissecting the adults using 26G needles. 2-cell embryos were quickly collected and transferred via mouth pipet to a 0.2ml PCR tube or to a poly-lysine coated slide on ice. Once 10-30 2-cell embryos were collected, the tube was placed in a BioRad PCR machine, pre-cooled to 15°C or the slide was placed into a humidified chamber at 15°C. Embryos were incubated at 15°C until the appropriate stage was reached, then shifted to 26°C to inactivate *zen-4*. After the inactivation period was over, embryos were shifted back to 15°C until the 1.5-2fold stage, when embryos from tubes were transferred to a 4% agar pad on a glass slide to assess foregut attachment by DIC microscopy or the slides were processed for immunostaining. Incubation times were determined based on Sulston’s embryonic lineage timing at 20°C, and adjusted accordingly; 15°C was considered 1.5x slower than 20°C, while 26°C was considered 1.2x faster. Timing for specific stages for N2 and *zen-4(or153ts)* was determined with test incubations (data not shown). The specific programs for each shift as follows:

#### Early shift (circa Arcade Cell birth)

360 minutes at 15°C (equivalent of 240 minutes at 20°C), 70 minutes at 26°C (equivalent of 84 minutes at 20°C), 270 minutes at 15°C (equivalent to 180 minutes at 20°C).

#### Mid shift (between birth and polarization)

480 minutes at 15°C (equivalent of 320 minutes at 20°C), 41 minutes at 26°C (equivalent of 49.2 minutes at 20°C), 140 minutes at 15°C (equivalent to 93 minutes at 20°C).

#### Late shift (circa polarization)

540 minutes at 15°C (equivalent of 360 minutes at 20°C), 70 minutes at 26°C (equivalent of 84 minutes at 20°C), 60 minutes at 15°C (equivalent to 50 minutes at 20°C).

TS shifts for epidermal cells were performed in a similar manner, in which dissected 2-cell embryos were transferred to a poly-lysine coated slide and incubated in a humidified chamber at the appropriate temperature. Control embryos were incubated at 15°C for 525 minutes (equivalent of 350 minutes at 20°C). Shifted embryos were incubated for 195 minutes at 15°C (equivalent of 130 minutes at 20°C), 70 minutes at 26°C (equivalent of 84 minutes at 20°C), 205 minutes at 15°C (equivalent to 137 minutes at 20°C).

Embryos for arcade cell shifts were scored for unattached foreguts using DIC optics, on a Zeiss Axioimager M2 upright compound microscope with a 63x objective. Embryos for epidermal experiments were fixed and stained and assessed for multinucleate cells and polarity using a Zeiss LSM880 confocal microscope.

### Molecular biology

A complete list of the primers used can be found in Table S6. For all PCR reactions, either AccuStart HiFi Taq Polymerase (QuantaBio) or PrimeStar Taq Polymerase (Takara) was used.

pML902 (*Pdlg-7kb*::DLG-1::GFP::*unc-54* 3’UTR) (McMahon *et al.*, 2001) was used as the template to generate *Pdlg-3.9*::DLG-1::GFP::*unc-54* 3’UTR and *Pdlg-4.9*::DLG-1::GFP::*unc-54* 3’UTR. After amplification, a Qiagen PCR Purification kit was used to clean up the reaction. The PCR products were subsequently used for injection to generate transgenics (see below).

ZEN-4::GFP (bsem1129) was generated by swapping out GFP for YFP from bsem1105 (ZEN-4::YFP) using NcoI and BamHI. bsem1129 contains genomic DNA from 3kb upstream of the predicted start site, the entire coding region up to the stop codon, fused with GFP and the *unc-54* 3’UTR from pPD95.77 (a gift of Andy Fire, Addgene plasmid # 1495).

ZEN-4 deletion constructs were generated by a variation of overlap PCR (Hobert, 2002), where the overlap primers contained sequence adjacent to the region to be removed and replaced with a KasI restriction site. The specific nucleotides (and the corresponding amino acids) removed in each construct can be found in Supplemental Data. Nuclear localization sites were predicted by PSORTII (http://psort.hgc.jp/form2.html) (Nakai and Horton, 1999). bsem1129 was used as the template to generate each 5’ and 3’ fragment, which were cleaned using the Qiagen PCR purification kit. 10ng of each 5’ and 3’ fragment were used for the overlap reaction. Overlap products were gel-purified (Qiagen) and cloned into pGEM-T (Promega) or pCR4Blunt-TOPO (Invitrogen). The NLS1/NLS2 double deletion was generated by subcloning a BclI-generated fragment from bsem1231 (ZEN-4::NLS1Δ::GFP) into the BclI site of bsem1210 (ZEN-4::NLS2Δ::GFP). Constructs were verified to have the expected nucleotides removed, and no additional mutations, by Sanger sequencing.

CYK-4::GFP (bsem1289) was generated via overlap PCR by amplifying the *C. briggsae* CYK-4 genomic coding sequence (up to the stop codon) from bsem1093 and fusing it to GFP::*unc-54* 3’UTR from pPD95.77. AIR-2::GFP (bsem1292) was generated via overlap PCR by amplifying the AIR-2 genomic coding sequence (up to the stop codon) from N2 genomic DNA and fusing it to GFP::*unc-54* 3’UTR from pPD95.77. The resulting overlap PCR products were subcloned into pCR4-TOPOBlunt and sequence verified. Each half of the ZEN-4 tether constructs was generated by PCR, with the added tether sequence as an overhang. The flexible tether sequence chosen has successfully generated forced dimers in *Drosophila* (Neuhold and Wold, 1993; Castanon *et al.*, 2001).

*Pelt-7::*mCherry_pGEM-T (bsem1146), *Pdpy-13*::mCherry_pGEM-T (bsem1178), *Pelt-7*::mCherry::HIS-58/H2B_pGEM-T (bsem1177) were generated by overlap PCR. The 2.9kb *elt-7* promoter was amplified from wildtype genomic DNA and fused with either mCherry::*unc-54* 3’UTR from pCFJ90 (gift of Christian Frokjaer-Jansen/Erik Jorgensen, Addgene plasmid # 19327) (Frokjaer-Jensen *et al.*, 2008) or with mCherry::H2B::*unc-54* 3’UTR from bsem1145. bsem1145 was generated by releasing mCherry::HIS-58/H2B from *pie-1::*mCherry::HIS-58/H2B (gift of Jon Audhya) (McNally *et al.*, 2006) using BamHI (NEB) and subcloning this fragment into the BamHI site of bsem1176 (*unc-54*_3’UTR_TOPO). The 260bp *dpy-13* promoter was amplified from wildtype genomic DNA using primers with attB1 and attB2 overhangs. The resulting product was recombined in to pDONR221 using BP Clonase II (Invitrogen) to create bsem1184. The *dpy-13* promoter was then amplified from bsem1184 and fused with mCherry::H2B::*unc-54* 3’UTR amplified from bsem1145 The overlap products were then subcloned into pGEM-T (Promega).

### Microinjection/Transgenic generation

2ng of linearized ZEN-4::GFP or of the deletion constructs described above were injected into SM1052 (*zen-4(px47) dpy-20(e1282)/bli-6(sc16) unc-24(e138)*) along with 5-10ng of either linearized bsem1146, bsem1177, and/or bsem1178 (fluorescent co-selectable markers), 55ng linearized pRF4 (*rol-6(d)* co-selectable marker), and Salmon Testes DNA (amount necessary to bring total DNA in injection mix to 100ng; Sigma D1626) to generate complex array-containing Roller animals.

ZEN-4 tether constructs were generated by *in vivo* recombination in the nematode gonad. Each half of the tether was generated by PCR and injected in equimolar amounts along with bsem1177, bsem1178, pRF4 and Salmon Testes DNA.

1ng of *Pdlg-1-*containing purified PCR product (see above), 55ng linearized pRF4 (*rol-6(d)* co-selection marker), and 44ng Salmon Testes DNA (Sigma D1626) were injected into N2 (SM1880) to generate complex array-containing Roller animals.

### RNAi

RNAi was performed as described (Timmons *et al.*, 2001), with minor modifications. A single bacterial colony (either *mCherry* as a negative control or *pha-4*) was picked to grow in 6ml of Luria broth + antibiotic (25ug/ml kanamycin) for 8h at 37°C. Bacteria were pelleted and resuspended in an IPTG/antibiotic mixture (8mM IPTG, 25ug/ml kanamycin), and 40ul was spread onto a 35mm NGM (Nematode Growth Media (Stiernagle, 2006)) plate. The plates were covered with foil and incubated at room temperature for 48h before use. N2 gravid adults were bleached to release eggs, which were hatched overnight in M9 without food to synchronize the population. Starved L1s were transferred to OP50 containing NGM plates and grown to the L4 stage at 20°. L4s were transferred to a conical tube and washed in M9 2-3x to remove all traces of OP50 bacteria. ~80 L4s were dropped onto either an mCherry or a *pha-4* RNAi plate and incubated at 25°C for 24h. The now adults were transferred to a new RNAi plate and incubated for an additional 6h at 25°C. Laid embryos were collected and either incubated overnight at 25°C on NGM plates to score terminal phenotypes or fixed with PFA immediately for immunostaining.

### Live Imaging

Embryos were placed on a 4% agar pad (Difco Noble Agar) in 10mM levamisole. A #1.5 coverslip (Corning) was added, and the nail polish was added at the corners to prevent the coverslip from moving. Embryos were scored using Nomarski optics for foregut attachment defects under a Zeiss AxioImager M2 running AxioVision software. For live-imaging of GFP fluorescence, embryos were either imaged under an AxioImager M2 with Apotome to remove out of focus light, or under the Zeiss LSM710, LSM780 or LSM880 confocal microscope.

### Immunostaining

Embryos were placed on poly-L-lysine (Sigma P-8920) coated slides in 50µl of water and processed using the freeze-crack method (Duerr, 2006; Shakes *et al.*, 2012). Following cracking, slides were incubated in ice-cold MeOH for 8 min. The exception was for *pha-4(RNAi)* processed embryos, where the water was removed and replaced by PFA fix (2% PFA diluted from a 16% methanol-free stock (Alfa Aesar) in 1x PBS) and then incubated for 3 min in ice-cold MeOH after cracking. Slides were blocked in TNB Buffer (100mM Tris-HCl pH 7.5, 200mM NaCl, 1% BSA) with 10% normal goat serum (NGS; VWR

#102643-580). All antibodies were also diluted using TNB + NGS. The following antibodies were used: mouse anti-ERM-1 (1:10, DSHB) (Hadwiger *et al.*, 2010); mouse anti-PAR-3 (1:10, DSHB P4A1) (Nance *et al.*, 2003); rabbit anti-PHA-4 (1:1000) (Kaltenbach *et al.*, 2005); mouse anti-PSD95 (detects DLG-1; Pierce MA1-045; 1:250) (Firestein and Rongo, 2001); rabbit anti-PAR-6 (1:1000, gift of Tony Hyman, Max Planck Institute) (Hoege *et al.*, 2010); and chicken anti-GFP (1:200, Millipore # AB16901). As noted, ERM-1 antibodies were obtained from the Developmental Studies Hybridoma Bank (DSHB) developed under the auspices of the NICHD and maintained by The University of Iowa, Department of Biology, Iowa City, IA 52242. Secondary antibodies conjugated to Alexa488, Alexa568 or Alexa647 were obtained from Invitrogen. Images were obtained on a Zeiss LSM880 confocal microscope, images were exported from ZEN Blue (Zeiss), maximum projections generated in ZEN Black (Zeiss), and rotated, resized and cropped in Adobe Photoshop.

### smFISH procedure and data processing

smFISH was performed mostly as described (Raj and Tyagi, 2010), and summarized here with some modifications. Custom Stellaris FISH Probes were designed against *dlg-1, erm-1, par-6*, and *gfp* by utilizing the Stellaris FISH Probe Designer (Biosearch Technologies, Inc., Petaluma, CA) available online at http://www.biosearchtech.com/stellarisdesigner. Exact probe sequences available in Supplemental Materials. *dlg-1, erm-1* and *par-6* probes were labeled with Quasar 570 dye, and *gfp* was labeled with Quasar 670. Probes were resuspended in 400ml TE (10mM Tris pH 8.0, 1mM EDTA pH 8.0), aliquoted, and stored at −20°C. Probes in use were stored at 4°C. One to five 60mm NGM plates with gravid adults and many laid embryos were washed with water to remove adults and larvae into a 15ml conical tube. To release the laid embryos, additional water was added to the plate and lightly rubbed with a gloved fingertip. After 1-2 washes in ddH_2_O, pelleted embryos and worms were transferred to a 1.5ml centrifuge tube and pelleted. To release the young embryos from the gravid adults, 100-200ul bleach solution (240ul bleach, 50ul 5M NaOH, 710ul ddH_2_O) was added to the pellet and incubated for 5 minutes at RT using a thermomixer (Eppendorf) to mix the sample every 30s. The embryos were quickly washed 3x using M9 buffer. The pelleted embryos were then fixed in 100-200ul 3.7% formaldehyde solution (100ul 37% Formaldehyde, 100ul 10x PBS, 800ul ddH_2_O) for 15 minutes at RT using a thermomixer to mix the sample every 3 minutes. The tube was flash-frozen in liquid nitrogen for 1 minute to freeze-crack the eggshells, then either stored at −80°C or processed immediately. The frozen sample was thawed at RT, then washed 2x with 200ul 1X PBS. The pelleted embryos were resuspended in 200ul 70% EtOH and stored overnight at 4°C. The next day the EtOH was exchanged for wash buffer (0.5ml formamide, 0.5ml 20x SSC (Invitrogen), 4ml ddH_2_O), then the sample was pelleted. 50ul of hybridization buffer (100ul deionized formamide (Calbiochem OmniPur/EMD Millipore), 0.1g dextran sulfate (Acros), 100ul 20x SSC, 800ul ddH_2_O) containing smFISH probes were added to the embryos (0.75ul for *dlg-1* probes, 1ul for *erm-1* and *par-6* probes, and 2ul for *gfp* probes), and incubated for 4 hours at 37°C in a thermomixer, mixing every 15 minutes. The embryos were rinsed once quickly using wash buffer, then washed for 1h at 37°C (mixing every 5 minutes). Pelleted embryos were resuspend in 50-100ul ddH_2_O, then added to poly-lysine coated slides and allowed to settle. ddH_2_O was removed and 7ul of SloFade with DAPI was added. A #1.5 coverslip was added, excess mounting material was wicked away with a Kimwipe, and sealed with clear nail polish. Slides were stored at −20°C until ready to image. Images were obtained on a Zeiss LSM880 confocal microscope with a Plan-Apochromat 63x/1.4 Oil objective using either a 568 laser (Quasar 570 probes alone) and a 633 laser (Quasar 670). GFP protein expression survived the fixation method and could be visualized with a 488 laser. Selected z-stacks that encompassed the digestive tract but excluded the epidermis were exported from ZEN Blue (Zeiss) and imported into Imaris 3D software (http://www.bitplane.com) for converting the smFISH signal into painted spheres.

### PHA-4 and FoxA ChIP-Seq analysis

For the *C. elegans* analysis, bed files from the early and late embryo PHA-4 ChIP-Seq datasets (as well as the MACS2 peak analysis bed file) (Zhong *et al.*, 2010), and the Ahringer and Meyer transcription start site datasets (Chen *et al.*, 2013; Kruesi *et al.*, 2013) were downloaded and visualized in IGV. Genes known to be expressed in epithelial tissue (those listed in Table 1 and Table S1) were queried, and a gene was annotated as PHA-4 bound if a MACS2-called PHA-4 peak was found upstream, at the TSS or within the first intron.

For the mammalian analysis, FOXA1/FOXA2 ChIP-Seq data was downloaded as peak calls for the following datasets: FoxA2 ChIP-Seq from mouse liver (Soccio *et al.*, 2011); FOXA1 ChIP-Seq from HEPG2 cells (Motallebipour *et al.*, 2009); FOXA2 ChIP-Seq from Caco2 cells (Gosalia *et al.*, 2015); and FOXA1 and FOXA2 from hESC differentiated into gut tube or foregut cells (Wang *et al.*, 2015). For the Caco2 dataset, two replicates were downloaded and the intersection was used for subsequent analysis. For FOXA1 Chip-Seq from MCF7 cells (Hurtado *et al.*, 2011), two replicates in FASTQ format were obtained and then processed to call peaks. Quality trim of reads was performed using Trimmomatic (Bolger *et al.*, 2014), mapping of treatment replicates and control input to genome was performed using Bowtie2(http://bowtie-bio.sourceforge.net/bowtie2/index.shtml) (Langmead and Salzberg, 2012), and peaks were called using MACS2 (https://pypi.python.org/pypi/MACS2) (Zhang *et al.*, 2008). The intersection of the replicate peaks was generated and used for subsequent analysis. Where needed, the input files were lifted to either mouse mm10 (GRCm38) or to human hg38 (GRCh38) genomes using the UCSC LiftOver program (http://genome.ucsc.edu/cgi-bin/hgLiftOver). Mouse and human gene annotation as well as mouse and human orthologs of *C. elegans* genes were downloaded from Ensembl (release 80). FOXA peaks that were up to 1000bp upstream, up to 1000bp downstream, or within the gene body of mouse and human genes were identified, and the final output was homologs of the 50 *C. elegans* PHA-4 bound, epithelial-specific genes (Table 1) that had FOXA binding as per these parameters. A manual inspection was performed to determine the percent of genes that had FOXA binding upstream or within the first or second intron. An affirmative call did not exclude binding in other areas beyond upstream or introns 1-2.

## ACKNOWLEDGEMENTS

We thank J. Nance for *par-6(MZ)* reagents; the Developmental Studies Hybridoma Bank (DSHB) and T. Hyman for antibodies; M. Labouesse for DLG-1::GFP plasmids and worm strains; E. Jorgensen, C. Frokjaer-Jansen, and J. Audhya for plasmids; the Harvard Center for Biological Imaging (HCBI) for help optimizing smFISH imaging; and J. Lisack and L. Rosen for technical help. Some strains were obtained from the CGC, which is funded by NIH P40 OD010440. Funding was supplied by Harvard University, the University of Utah, the MacArthur Foundation, NIH 5F32 GM084650 (SEV), NIH 1R21 HD066263 (SEM), and NIH 5R37 GM056264 (SEM).

## Abbreviations

ZEN: Zygotic ENclosure defective
PAR: PARtitioning defective
PHA: defective PHArynx development
ERM: Ezrin/Radixin/Moesin homolog
DLG: Disks LarGe
CYK: CytoKinesis defective
Crb: Crumbs
Scrib: Scribble
LGL: Lethal Giant Larvae
FOXA: Forkhead bOX transcription factor subgroup A
smFISH: single molecule Fluorescent In Situ Hybridization
ChIP-Seq: Chromatin ImmunoPrecipitation-Sequencing
TSS: Transcription Start Site
MET: Mesenchymal-to-Epithelial Transition
EMT: Epithelial-to-Mesenchymal Transition

